# Cadherin-dependent adhesion is required for muscle stem cell niche anchorage and maintenance

**DOI:** 10.1101/2023.10.05.561107

**Authors:** Margaret Hung, Hsiao-Fan Lo, Aviva G. Beckmann, Deniz Demircioglu, Gargi Damle, Dan Hasson, Glenn L. Radice, Robert S. Krauss

## Abstract

Adhesion between stem cells and their niche provides stable anchorage and signaling cues to sustain properties such as quiescence. Skeletal muscle stem cells (MuSCs) directly adhere to an adjacent myofiber via cadherin-catenin complexes. Previous studies on N- and M-cadherin function in MuSCs revealed that while N-cadherin is required for quiescence, they are collectively dispensable for MuSC niche localization and regenerative activity. While additional cadherins are expressed at low levels, these findings raise the possibility that cadherins are unnecessary for MuSC anchorage to the niche. To address this question, we conditionally removed from MuSCs β- and γ-catenin and, separately, αE- and αT-catenin, factors essential for cadherin-dependent adhesion. Catenin-deficient MuSCs break quiescence similarly to N-/M-cadherin-deficient MuSCs, but exit the niche, and are depleted. A combination of in vivo, ex vivo, and single cell RNA sequencing approaches reveal that MuSC attrition occurs via a single fate: precocious differentiation, reentry to the niche, and fusion to myofibers. These findings indicate that cadherin-catenin–dependent adhesion is required for anchorage of MuSCs to their niche and preservation of the stem cell compartment. Furthermore, separable, cadherin-regulated functions govern niche localization, quiescence, and stem cell maintenance in MuSCs.

**SUMMARY STATEMENT:** Genetic ablation of cadherin-based adhesion in skeletal muscle stem cells triggers activation, niche exit, precocious differentiation, and subsequent depletion of the stem cell pool.

## INTRODUCTION

Adult stem cells reside in a specialized microenvironment, or niche, that supplies signals to sustain specific cellular properties (Fuchs & Blau, 2020). A general feature provided by the niche is anchorage of stem cells via cell-cell and/or cell-extracellular matrix adhesion (Chen et al., 2013; Parsons et al., 2010; Polisetti et al., 2016; Schüler et al., 2022). Niche-based adhesion in turn supports stable proximity of stem cells to biochemical and biomechanical signals, the nature of which may vary between tissues and stem cell types. Skeletal muscle stem cells (MuSCs, also called satellite cells) are localized between a myofiber and an enwrapping basal lamina. During homeostasis MuSCs are quiescent, but in response to muscle injury they activate, proliferate, differentiate, and fuse with each other or existing myofibers to repair the damage (Hardy et al., 2016; Schmidt et al., 2019). They also self-renew and reoccupy the niche in regenerated muscle.

Niche localization induces polarity in MuSCs; laminin-binding integrins are found basally and cadherins are present apically, the latter being the cell surface where MuSCs are in direct contact with myofibers (Goel et al., 2017; Krauss et al., 2017; Rozo et al., 2016). α7β1-integrin is the major laminin receptor in MuSCs, and conditional genetic removal of β1-integrin from these cells results in loss of cell polarity and eventual MuSC attrition due to precocious differentiation (Rozo et al., 2016). Despite this perturbance, β1-integrin-deficient MuSCs remain under the basal lamina. Therefore, β1-integrin-mediated basal adhesion is required for maintenance of the MuSC compartment, but additional mechanisms are sufficient to retain MuSCs in the niche.

The roles of apical adhesion and cadherins in MuSCs are less clear. Cadherins are calcium-dependent, homophilic cell-cell adhesion molecules that regulate tissue integrity and specific morphogenetic movements (Yap et al., 2018; Mège & Ishiyama, 2017). Classical cadherins have a cytoplasmic domain bound directly by β- and/or γ-catenin, which in turn bind to α-catenin. α-catenin binds directly to F-actin or to adaptor proteins that bind F-actin, forming molecular complexes that provide stable cell-cell cohesion and transmit tensile forces from the exterior to the interior of the cell (Yap et al., 2018). MuSCs express multiple catenin-binding cadherins, including M-cadherin, N-cadherin, and VE-cadherin, each localized to the MuSC apical membrane (Fukada et al., 2007; Goel et al., 2017). M-cadherin (encoded by *Cdh15*) expression is highly enriched in the skeletal muscle lineage, and *Cdh15* mRNA is expressed at much higher levels in quiescent MuSCs than *Cdh2* or *Cdh5* mRNAs (encoding N-cadherin and VE-cadherin, respectively) (Yue et al., 2020). Surprisingly, germline mutation of *Cdh15* in mice did not result in an obvious phenotype in skeletal muscle development, MuSC number, or regeneration (Goel et al., 2017; Hollnagel et al., 2002). In contrast, conditional mutation of *Cdh2* in MuSCs rendered the cells prone to break quiescence and enter a state of partial activation, without compromising cell polarity or niche localization (Goel et al., 2017). Furthermore, MuSCs lacking *Cdh2* are regeneration- and self-renewal-proficient (Goel et al., 2017). Combined loss of *Cdh15* and *Cdh2* exacerbated the phenotypes associated with loss of *Cdh2* alone, but double-mutant cells also remained in the niche and are functional in response to injury (Goel et al., 2017). Therefore, the most abundantly expressed cadherin, M-cadherin, is dispensable for MuSC function, whereas N-cadherin is expressed at much lower levels, yet it is required to prevent MuSCs from entering the activation process in the absence of injury. N-cadherin’s role in maintaining MuSC quiescence is linked to its function in promoting outgrowth and/or maintenance of cellular projections that apparently act as sensors of the niche (Kann et al., 2022).

VE-cadherin is present at the apical membrane of MuSCs lacking M- and N-cadherin (Goel et al., 2017). Additionally, quiescent MuSCs express RNA for at least one additional catenin-binding cadherin, albeit at low levels (R-cadherin, encoded by *Cdh4*) (Yue et al., 2020). Despite the presence of cadherins other than M- and N-cadherin, these findings raise the possibility that any cadherin-based adhesion may be dispensable for anchorage of MuSCs to their niche. Integrin-mediated basal adhesion and/or alternative mechanisms of apical adhesion may be sufficient for niche localization, even in the absence of the most abundant cadherin (M-cadherin) and a cadherin required for quiescence (N-cadherin). To address these possibilities, we assessed the consequences of depriving MuSCs of all ability to form cadherin-based adhesion structures. Simultaneous genetic removal of all expressed catenin-binding cadherins would require removing at least four cadherins, so we used conditional mutagenesis to target catenins, essential components of cadherin-based adhesion complexes. We separately removed two pairs of redundant catenins from MuSCs: β- and γ-catenin, and αE- and αT-catenin. We report here that MuSCs lacking these catenin pairs, and thereby deprived of cadherin-based adhesion, spontaneously activate and exit the niche, entering the interstitial space between myofibers. These cells are lost over time to what appears to be a single fate: precocious differentiation, reentry to the niche, and fusion to myofibers. Therefore, cadherin-based cell-cell adhesion is required for niche localization and preservation of MuSCs, and cadherins expressed at low levels appear to be sufficient for this stem cell property. Furthermore, the requirement for cadherins in MuSC niche localization and quiescence are distinguishable.

## RESULTS

### Conditional deletion of α-catenins or β- and γ-catenin results in MuSC attrition

Cadherins require β- or γ-catenin plus α-catenin to form stable cell adhesion junctions (Mège & Ishiyama, 2017). To determine the effects of loss of cadherin-based adhesion complexes, conditional mutagenesis approaches were taken. MuSCs express two isoforms of α-catenin, αE and αT, respectively encoded by *Ctnna1* and *Ctnna3* (Yue et al., 2020). Mice homozygous for floxed alleles for both genes were bred to mice that carried a MuSC-specific *Pax7^CreERT2^*allele, yielding *Ctnna1^fl/fl^;Ctnna3^fl/fl^;Pax7^CreERT2^*mice (referred to as α-cdKO mice; Fig. 1A). MuSCs express both β- and γ-catenin, encoded by *Ctnnb1* and *Jup*, respectively (Goel et al., 2017; Yue et al., 2020). A similar approach was taken to produce *Ctnnb1^fl/fl^;Jup^fl/fl^;Pax7^CreERT2^*mice (referred to as βγ-cdKO mice; Fig. 1G). Two-to-three month old mice were treated with tamoxifen (TMX) for five days to activate Cre and studied at various time points afterwards for catenin protein expression and MuSC homeostasis as assessed by number of Pax7^+^ cells.

**Fig. 1.**
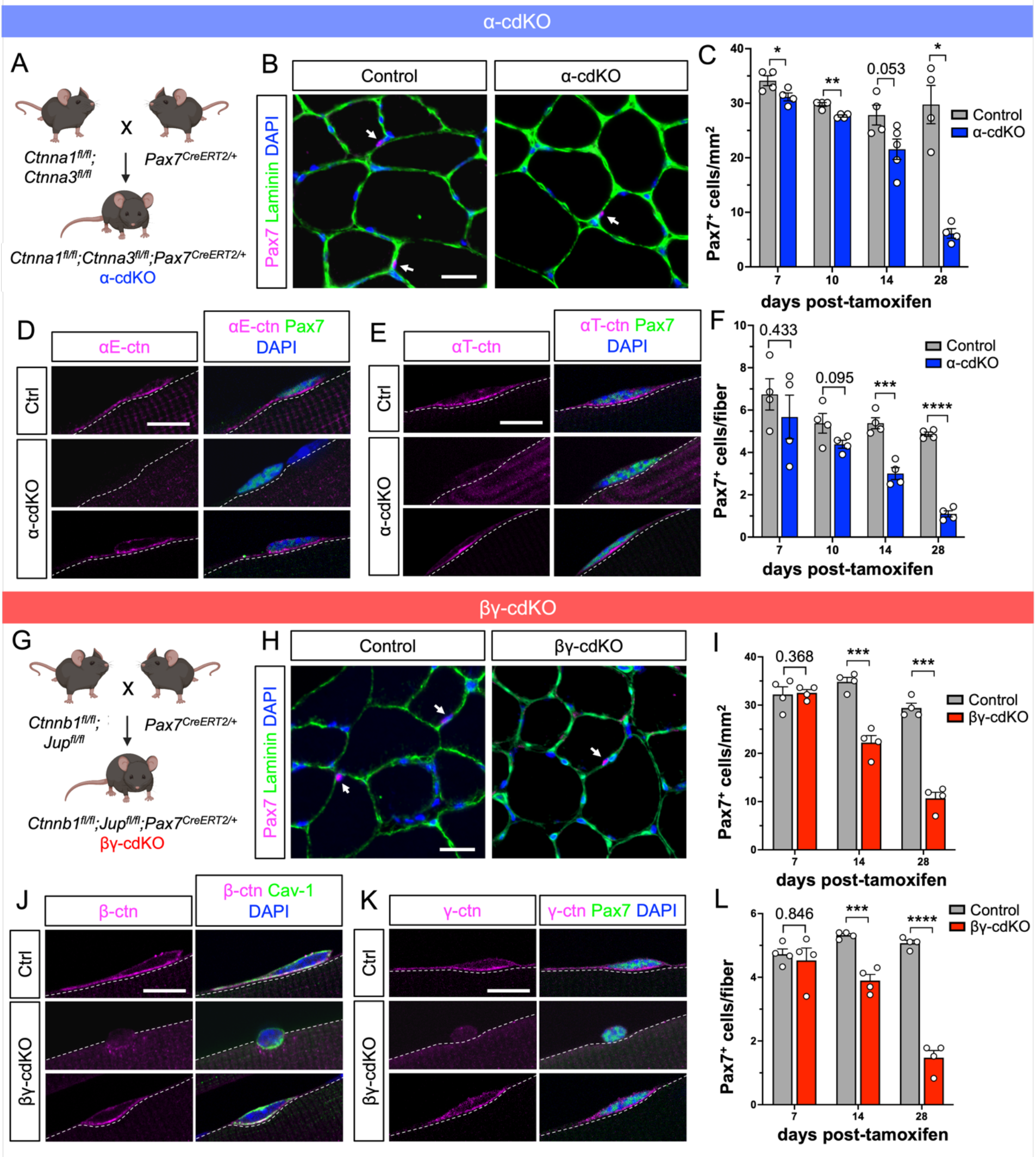
Catenins are apically enriched in quiescent MuSCs and necessary for MuSC maintenance. **(A)** Breeding scheme for α-cdKO mice: mice carrying homozygous floxed alleles for αE-catenin (*Ctnna1*) and αT-catenin (*Ctnna3*) loci were paired with mice carrying a MuSC-specific Pax7-driven CreERT2 (Pax7^CreERT2^) allele to yield α-cdKO mice. Made with BioRender. **(B-F)** TA muscle sections from control and α-cdKO mice at various time points were immunostained (Pax7, magenta; laminin, green; DAPI, blue) and Pax7-expressing MuSCs were labeled (B) and quantified (C). Immunostaining of single myofibers (αE- or αT-catenin, magenta; Caveolin-1 or Pax7, green; DAPI, blue) was used to assess presence of αE- and αT-catenin protein (D,E) and MuSC attrition (F) in α-cdKO mice. **(G)** Breeding scheme for βγ-cdKO mice: mice carrying homozygous floxed alleles for β-catenin (*Ctnnb1*) and γ-catenin (*Jup*) loci were paired with mice carrying a MuSC-specific Pax7-driven CreERT2 allele (Pax7^CreERT2^) to yield βγ-cdKO mice. Made with BioRender. **(H-L)** TA muscle sections from control and βγ-cdKO mice at various time points were immunostained (Pax7, magenta; laminin, green; DAPI, blue) and Pax7-expressing MuSCs were labeled (H) and quantified (I). Immunostaining of single myofibers (β- or γ-catenin, magenta; Pax7, green; DAPI, blue) was used to assess presence of β- and γ-catenin protein (J,K) and MuSC attrition (L) in βγ-cdKO mice. Scale bars: 25 μm (B,H), 10 μm (D,E,J,K). Each data point represents the average from ten fields (TA muscle sections) or at least ten myofibers (EDL single myofibers) from each animal. Data represent n≥4 per genotype per timepoint and represented as mean ± S.E.M with comparisons by two-tailed unpaired Student’s t-test. * = p < 0.05, ** = p < 0.01, *** = p < 0.001, **** = p < 0.0001.

α-cdKO mice displayed a steady decline in the number of Pax7^+^ MuSCs beginning approximately 10 days after the last TMX injection. This was observed in cross-sections of tibialis anterior (TA) muscle over a 28-day time course, at which point α-cdKO mice had lost 78% of Pax7^+^ MuSCs as compared to their control counterparts (Fig. 1B,C). Very similar results were obtained when quantifying Pax7^+^ MuSCs on single myofibers prepared from extensor digitorum longus (EDL) muscles (Fig. 1D-F). Single myofiber preparations were also used to assess expression of αE- and αT-catenin by immunofluorescence. Control MuSCs were uniformly positive for αE- and αT-catenin proteins, which were enriched at the apical membrane of each cell. Of the 22% of MuSCs remaining on EDL myofibers at 28 days post-TMX (DPT), 82% were αE-catenin immunoreactive and 78% were αT-catenin immunoreactive, while the rest were negative (Fig. 1D,E). The fraction of MuSCs that remained α-catenin^+^ had very likely failed to recombine all four floxed alleles, or some sufficient number of alleles, to cause cell loss.

We performed parallel analyses with βγ-cdKO mice. The phenotype of βγ-cdKO mice at 28 DPT was similar to that of α-cdKO mice, with loss of 64% and 71% of Pax7^+^ MuSCs in cross-section analyses of TA muscle and on single EDL myofibers, respectively (Fig. 1H,I,L). Of the 29% of MuSCs remaining on EDL myofibers, 53% were β-catenin^+^ and 66% were γ-catenin^+^, while the rest were negative (Fig. 1J,K). Again, the fraction of MuSCs remaining β- and/or γ-catenin^+^ at 28 DPT had likely failed to recombine all four floxed alleles, leaving enough residual protein to maintain MuSC polarity.

β-catenin plays critical roles in both cadherin-based adhesion and canonical Wnt signaling (van der Wal & van Amerongen, 2020). γ-catenin has overlapping function with β-catenin in binding classical cadherins but is not involved in Wnt signaling; it is however critical for formation of desmosomes, another type of cell-cell junction comprising desmosomal cadherins (Green et al., 2019; Kowalczyk & Green, 2013). β-catenin has been conditionally removed from adult MuSCs in previous reports, and it did not lead to MuSC attrition at homeostasis (Murphy et al., 2014; Rudolf et al., 2016). Additionally, quiescent MuSCs did not display canonical Wnt signaling activity. These results suggest that the phenotype of βγ-cdKO mice is due to loss of cell adhesion complexes, not due to perturbance of Wnt signaling. To provide additional evidence for this contention, we studied mice lacking only γ-catenin in MuSCs (γ-cKO mice). In contrast to βγ-cdKO mice, γ-cKO mice did not display any loss of Pax7^+^ MuSCs at 28 DPT, despite 94% of MuSCs on single myofibers lacking γ-catenin protein expression (Fig. S1). These results indicate that removal of both β- and γ-catenins is required for MuSC attrition. Combined with the similar phenotype, over a similar time course, seen in α-cdKO mice, our findings strongly suggest that loss of cadherin-based cell-cell junctions, rather than other functions provided by the various catenins, is the likely major cause of MuSC attrition in these mice.

### α-cdKO and βγ-cdKO mice display incomplete muscle regeneration and do not efficiently reoccupy the niche

MuSCs are essential for muscle regeneration (Lepper et al., 2011; Murphy et al., 2011; Sambasivan et al., 2011), but mice harboring only 10-20% of the total MuSC pool are still capable of muscle injury repair, indicating that regeneration is not dependent on large numbers of MuSCs (von Maltzahn et al., 2013). To assess whether the reduced MuSC populations in α-cdKO and βγ-cdKO mice were capable of regenerating muscle, we performed two consecutive BaCl_2_ injuries (Fig. 2A). Mice were maintained on TMX chow to continue Cre recombination stimulus during the regeneration period, and regeneration was assessed in both lines at 7 and 28 days post-injury (dpi) to observe early and complete regeneration, respectively.

**Fig. 2.**
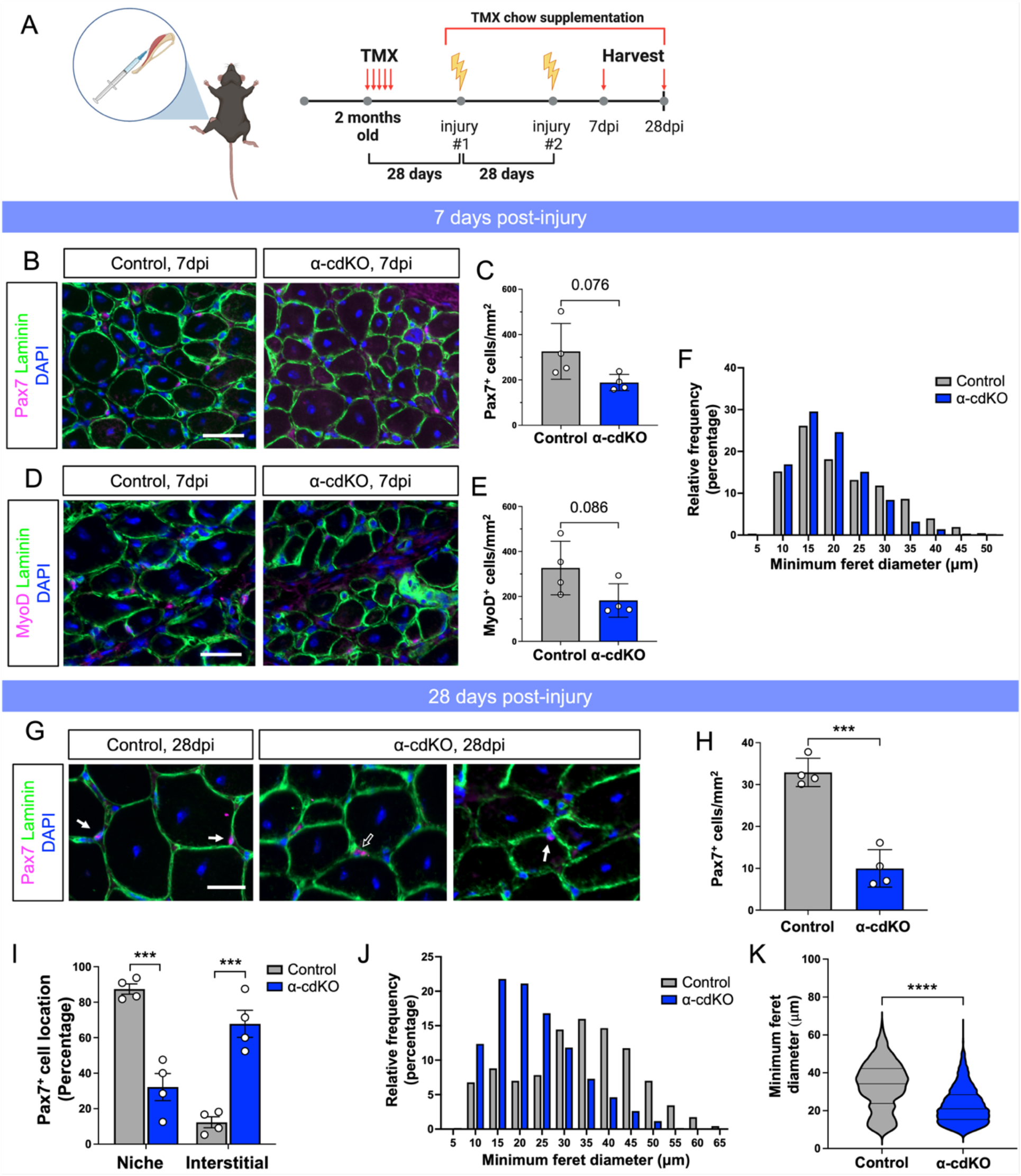
α-cdKO mice have modest defects in muscle regeneration. **A)** Double BaCl_2_ injury model. α-cdKO mice were injected with tamoxifen for 5 consecutive days to recombine α-catenin alleles. At 28 DPT, mice were placed on tamoxifen-supplemented chow, and the first injury was applied to the hindlimb by an intramuscular injection of 1.2% BaCl_2_ in saline. After 28 days of recovery, a second injury was applied. Muscle was harvested and snap frozen at 7 and 28 dpi for analysis. Made with BioRender. **(B-F)** At 7dpi, TA muscle sections from control and α-cdKO mice were immunostained (Pax7 or MyoD, magenta; laminin, green; DAPI, blue) for Pax7^+^ (B,C) and MyoD^+^ (D,E) cells to assess MuSC and myogenic progenitor numbers during early regeneration. Myofiber size was quantified by minimum feret diameter in (F). **(G-K)** At 28dpi, TA muscle sections from control and α-cdKO mice were immunostained (Pax7, magenta; laminin, green; DAPI, blue) to assess MuSC numbers (H) and location (I) after complete regeneration. Myofiber size was quantified by minimum feret diameter (J) with a statistical analysis in median myofiber size by two-tailed Mann-Whitney test (K). Scale bars: 25 μm (B,D,G). Solid arrow denotes cell under the basal lamina; open arrow denotes interstitial cell. Each data point represents the average from ten fields from each animal. All data represent n=4 per genotype and represented as mean ± S.E.M with comparisons by two-tailed unpaired Student’s t-test, unless otherwise noted. *** = p < 0.001, **** = p < 0.0001.

In α-cdKO mice, numbers of both Pax7^+^ MuSCs and MyoD^+^ myogenic precursor cells trended downward at 7dpi when compared to control mice, but fell short of p < 0.05 (Fig. 2B-E), which was closely mirrored in βγ-cdKO mice (Fig. S2A-D). Similarly, there was a slight shift toward smaller myofiber minimum feret diameter measurements in mutants (Figs. 2F, S2E). At 28dpi, the number of Pax7^+^ MuSCs in α-cdKO and βγ-cdKO mice had returned to a similar density to that at 28 DPT in uninjured mice (Fig. 1C,I), a ∼70% decrease relative to control mice (Figs. 2G,H, S2F,G). These data suggest that the initial, diminished pool of Pax7^+^ MuSCs in α-cdKO and βγ-cdKO mice had self-renewed but did not expand to refill the normal complement of stem cells. In contrast to what was observed at 7dpi, myofiber minimum feret diameter was strongly shifted towards smaller sizes at 28dpi (Figs. 2J,K, S2H), indicating a suboptimal regeneration response. Furthermore, 68% of α-cdKO MuSCs were localized in the interstitial space at 28dpi, compared to 12% in control MuSCs (Fig. 2G,I). This phenomenon was also observed in βγ-cdKO mice; 44% of βγ-cdKO MuSCs were found in the interstitial space, compared to only 13% of control MuSCs (Fig. S2F,H). Wild-type levels of catenin expression are therefore required for MuSCs to reoccupy the niche after injury. Regenerated muscles in α-cdKO and βγ-cdKO mice also displayed an increase in collagen deposition as detected by Picrosirius Red staining. This may be due to the presence of an increased number of smaller myofibers, thereby increasing circumferential fiber surface area per field, rather a primary regenerative defect (Figs. S2I,J, S3).

### α-cdKO MuSCs do not adopt non-myogenic fates

Initial homeostatic characterization and injury studies in α-cdKO and βγ-cdKO mice resulted in strikingly similar phenotypes, implicating loss of cadherin-catenin-based adhesion as an initiating event of MuSC attrition. To identify the fate of MuSCs lacking catenins during homeostasis, we focused on α-cdKO mice as a higher percentage of cells were affected following TMX treatment than in βγ-cdKO mice. We performed single cell RNA sequencing on purified α-cdKO MuSCs at 14 DPT, a timepoint midway into the process of MuSC attrition. A tdTomato reporter allele was crossed onto the α-cdKO line, and hindlimb muscles from three-month old control and α-cdKO mice were dissociated into single cell suspensions for FACS isolation of tdTomato^+^ cells. This protocol avoided sorting MuSCs based on expression of cell surface markers, which could have changed upon loss of α-catenins. Importantly, any cells that lost myogenic identity would still be captured for analysis.

Overall quality metrics assessed via CellRanger confirmed sample robustness before Harmony integration using all samples followed by unsupervised clustering of cells and non-linear dimensionality reduction via uniform manifold approximation and projection (UMAP) (Fig. S4A,B) (Korsunsky et al., 2019; McInnes et al., 2018). Eight unsupervised clusters were identified, none of which were unique to control or α-cdKO MuSCs (Fig. S4A,B). Following integration, cells were filtered for MuSC and myogenic progeny identity using UniCell Deconvolve unsupervised cell type annotations, with the great majority of cells identified as skeletal muscle stem cells or myogenic precursors (94%, 27,128 out of 28,790 total cells). Remaining cells excluded from further analyses were classified by UniCell as ‘striated muscle’ and excluded from downstream analyses (Fig. S4C) (Charytonowicz et al., 2023). Unsupervised clustering of MuSCs identified six clusters identified in both control and α-cdKO samples, showing similar characteristics to previously reported data sets (Figs. 3,S4E). A cluster of cells expressed transcripts for *Pax7*, *Gas1*, the Notch target *Hes1*, and receptor tyrosine kinase inhibitor *Spry1*, which revealed the isolation of a population similar to quiescent MuSCs *in vivo* (Cluster 2, Fig. 3A,B) (Dell’Orso et al., 2019; Machado et al., 2017). Many cells expressed the immediate early gene *Fos* within the same cluster, indicative of MuSCs in the earliest state of activation, likely due to the isolation process (Fig. 3A,C) (Almada et al., 2021; Kann et al., 2022; Machado et al., 2017; Van Velthoven et al., 2017). It is well established that the process of isolating and sorting MuSCs induces a stress response in, and activation of, MuSCs (Machado et al., 2017, 2021; Van Velthoven et al., 2017). We observed this phenomenon also (Fig. S4D). *Myod1* and *Myf5* expression was found in multiple clusters with varying degrees of expression and overlapped with *Pax7* expression and progressive MuSC activation (Clusters 0 and 1; Fig. 3A,C). In contrast, *Myog* expression, found only in cells committed to differentiate, was found in a single cluster (Cluster 4); these cells comprise a greater fraction in α-cdKO cells than control cells (p = 0.0434) (Fig. 3D). Overall, the changes in gene expression associated with MuSC isolation made it difficult to identify changes in early activation that occurred in α-cdKO MuSCs. However, this analysis of tdTomato-marked cells demonstrated that α-cdKO MuSCs did not adopt non-myogenic cell fates, indicating that they were lost by an alternative mechanism.

**Fig. 3.**
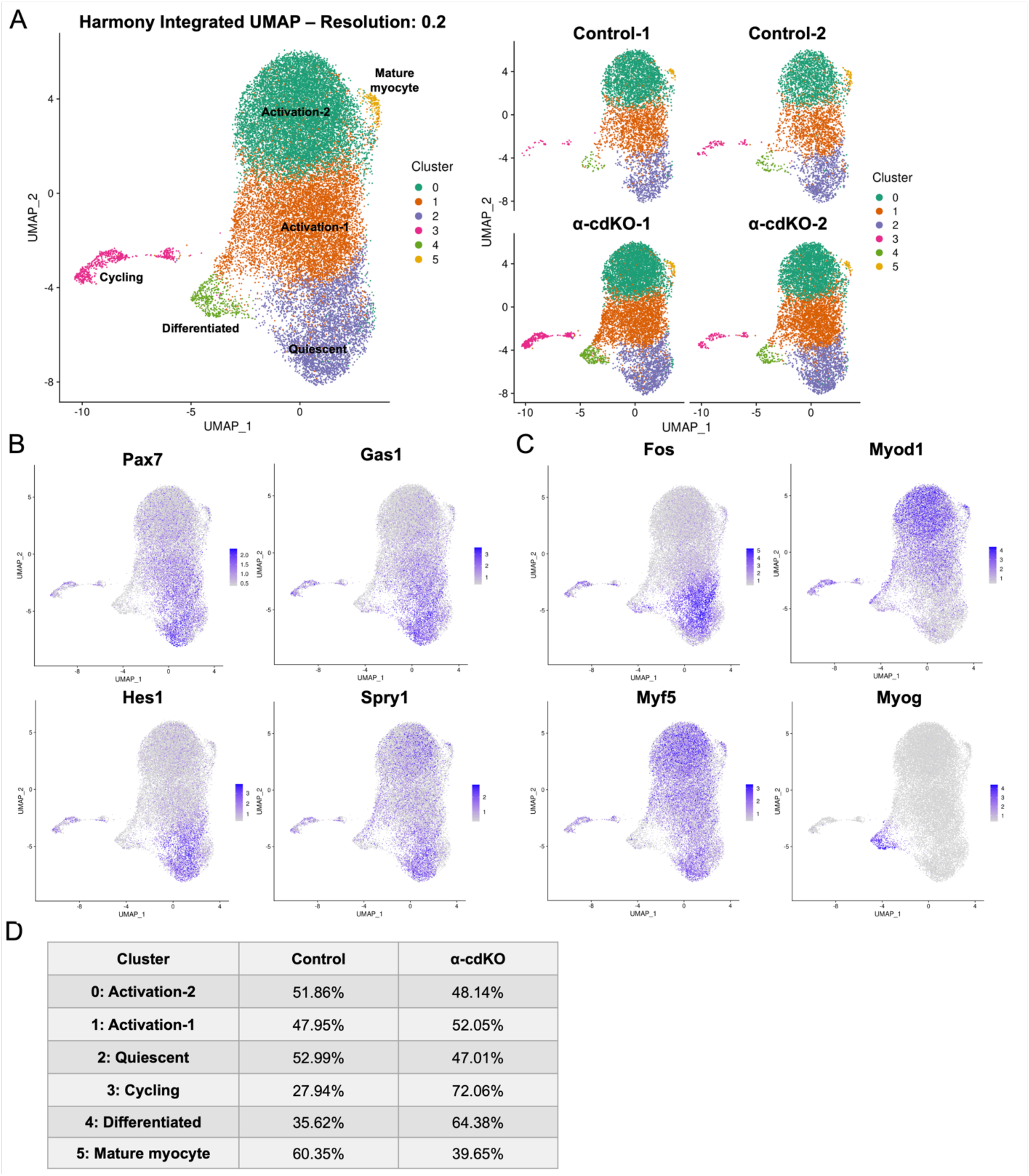
α-cdKO MuSCs do not adopt non-myogenic lineage fates. **(A)** Single cell RNA sequencing analysis of tdTomato^+^ cells from control and α-cdKO mice at 14 DPT revealed six distinct cell clusters after Harmony integration, with no cluster being unique to either genotype. **(B-D)** Gene expression of quiescence-associated (B) and activation- or differentiation-associated factors (C) illustrated a continuum of the quiescence-to-activation transition present in both genotypes. Clusters were categorized based on top differentially expressed genes, and genotype contribution to each cluster was normalized to total population and cluster population numbers (D).

### α-cdKO MuSCs undergo activation in the absence of injury

Quiescent MuSCs have long, heterogeneous projections and retraction of these structures is a very early response to muscle injury (Kann et al., 2022; Ma et al., 2022; Verma et al., 2018). Consistent with this observation, MuSCs characterized as being in a state of partial activation (e.g., *Cdh2*-mutant MuSCs and MuSCs in G_alert_) either lack or have shorter projections (Kann et al., 2022; Goel et al., 2017; Rodgers et al., 2014). To visualize and quantify MuSC projections we used α-cdKO mice carrying a tdTomato reporter and isolated single myofibers from these and control mice carrying the reporter at 14 DPT. α-cdKO MuSCs had fewer and shorter cellular projections than control MuSCs, consistent with the possibility of their having initiated the activation process in the absence of injury (Fig. 4A-C).

**Fig. 4.**
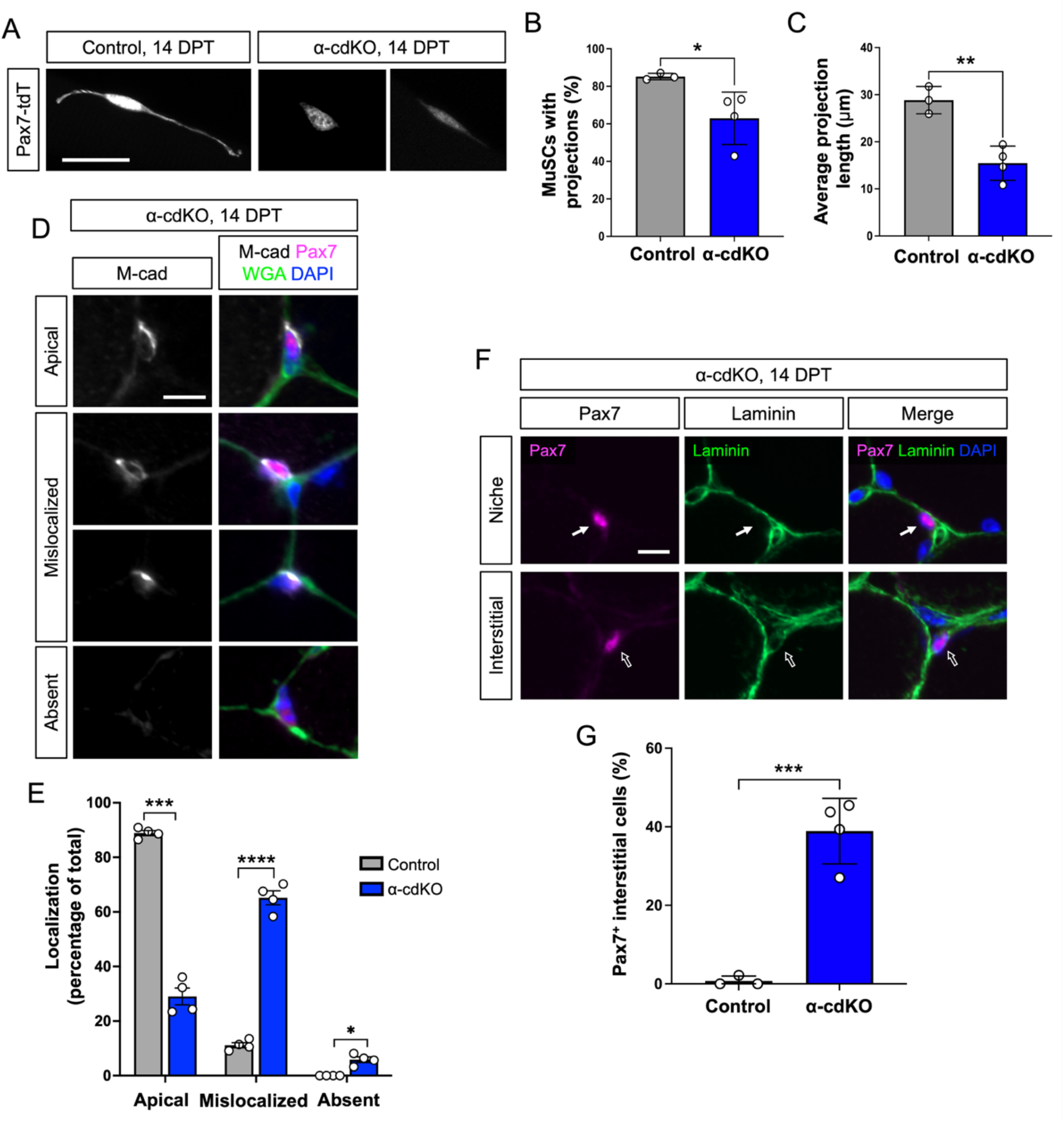
Loss of α-catenins compromises cadherin-based adhesion and leads to escape from the niche. **(A-C)** Endogenous tdTomato reporter fluorescence from MuSCs on isolated single EDL myofibers from control and α-cdKO mice at 14 DPT (A) was used to quantify the proportion of MuSCs maintaining cytoplasmic projections (B) and the average length of remaining projections (C). **(D-E)** At 14 DPT, TA muscle sections from control and α-cdKO mice at 14 DPT were immunostained (M-cadherin, grey; Pax7, magenta; WGA, green; DAPI, blue) to assess M-cadherin localization in Pax7^+^ MuSCs (D, E). **(F-G)** At 14 DPT, TA muscle sections from control and α-cdKO mice at 14 DPT were immunostained (Pax7, magenta; laminin, green; DAPI, blue) (F) to assess MuSC localization (G). Scale bars: 25 μm (A,F), 10 μm (D). Each data point represents the average quantified from n≥30 cells from each animal (TA muscle sections) or at least ten myofibers (EDL single myofibers) from each animal. Data represent n=4 per genotype and represented as mean ± S.E.M with comparisons by two-tailed unpaired Student’s t-test. * = p < 0.05, ** = p < 0.01, *** = p < 0.001, **** = p < 0.0001.

We next assessed cadherin protein distribution and niche localization. TA muscle sections were immunostained for Pax7 and M-cadherin, the latter serving as a readout for polarized distribution of cadherin-based junctions in MuSCs (Fig. 4D,E). In 89% of control MuSCs, M-cadherin was enriched at the apical membrane, the cell surface in direct contact with a myofiber. In contrast, 65% of α-cdKO MuSCs had M-cadherin present throughout the plasma membrane or enriched on the basal side of the cell. Additionally, 6% of mutant cells completely lacked M-cadherin signal. Therefore, in the absence of α-catenins, MuSCs were unable to maintain polarized apical localization of the major cadherin protein expressed in these cells. Consistent with loss of this characteristic property of MuSCs, 39% of Pax7^+^ α-cdKO MuSCs had exited the niche and were detected in the interstitial space between myofibers, while virtually all control MuSCs were found within the niche, under the basal lamina (Fig. 4F,G).

To further assess if α-cdKO MuSCs became activated and entered the cell cycle, TA sections were stained for Pax7 and the MuSC cell cycle and activation marker Ki67. While Pax7^+^ MuSCs in control mice rarely expressed Ki67, 35% of α-cdKO MuSCs were Ki67^+^ (Fig. 5A,B). Furthermore, 74% of Pax7^+^Ki67^+^ MuSCs were present in the interstitial space, outside the basal lamina (Fig. 5A,C). Approximately 77% of Ki67^+^ α-cdKO MuSCs were also positive for the activation and myogenic progenitor cell marker MyoD, with 97% of MyoD^+^Ki67^+^ cells present in the interstitial space (Fig. 5D-F). Taken together, α-cdKO MuSCs exhibited a range of phenotypes at 14 DPT, consistent with sequencing data (Fig. 3). While nearly all MyoD^+^Ki67^+^ cells had exited the niche, a fraction of α-cdKO Pax7^+^ MuSCs remained under the basal lamina and did not express Ki67 (Fig. 5A,B). Some of these latter cells may not have undergone complete loss of α-catenins, either due to incomplete Cre-mediated recombination or perdurance of a sufficient level of α-catenin protein to maintain cadherin-dependent adhesion. Alternatively, distinct, but not fully sufficient, adhesion mechanisms may exist leading to asynchronous MuSC activation and niche exit in the absence of α-catenins.

**Fig. 5.**
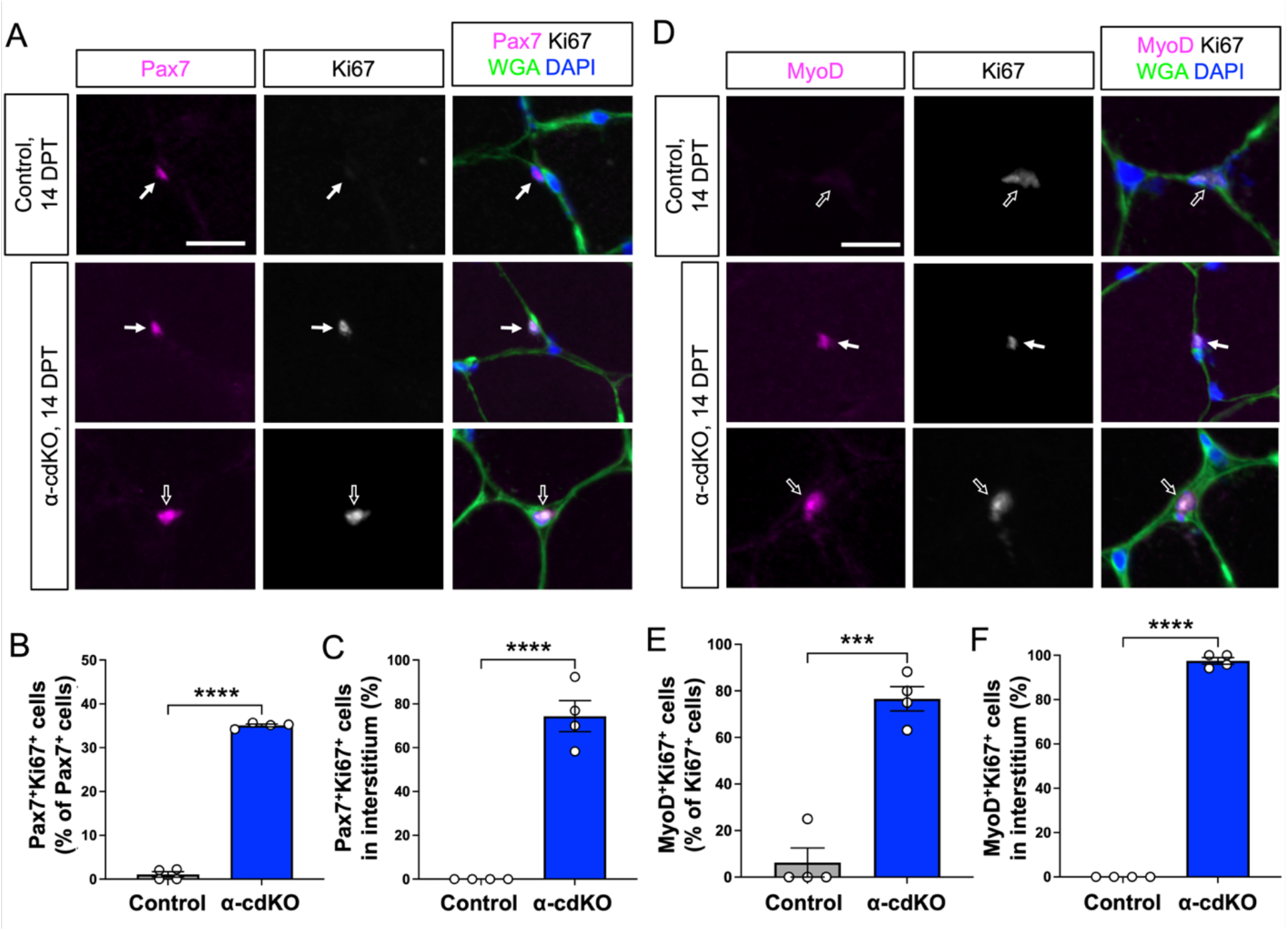
α-cdKO MuSCs activate and enter the cell cycle in the absence of injury. **(A-C)** TA muscle sections from control and α-cdKO mice at 14 DPT were immunostained (Pax7, magenta; Ki67, grey; WGA, green; DAPI, blue) (A) to assess incidence of Pax7^+^Ki67^+^ cells (B) and their location (C). **(D-F)** TA muscle sections from control and α-cdKO mice at 14 DPT were immunostained (MyoD, magenta; Ki67, grey; WGA, green; DAPI, blue) (D) to assess incidence of MyoD^+^Ki67^+^ cells among all Ki67^+^ cells (E) and their location (F). Scale bars: 10 μm (A,D). Each data point represents the average quantified from n≥30 cells from each animal. Solid arrow denotes cell under the basal lamina; open arrow denotes interstitial cell. Data represent n=4 per genotype and represented as mean ± S.E.M with comparisons by two-tailed unpaired Student’s t-test. *** = p < 0.001, **** = p < 0.0001.

### α-cdKO MuSCs progress through the myogenic lineage and are lost to precocious differentiation and fusion

A large fraction of α-cdKO MuSCs at 14 DPT displayed features of activated MuSCs. Attrition of these cells had begun by this point and continued for another 14 days (Fig. 1). We examined possible mechanisms by which α-cdKO MuSCs were lost, including apoptosis, and differentiation and fusion to myofibers. To assess apoptosis, TA muscle sections were stained for cleaved caspase-3 and analyzed by TUNEL assay, with sections of intestinal epithelium from the same animals as a positive control for the former and DNaseI-treated TA sections as a positive control for the latter. In control and α-cdKO muscles, cell death was not detected by either method (Fig. S5).

To validate the increased expression of *Myog* seen in our sequencing data and test for loss of α-cdKO MuSCs via differentiation and fusion with myofibers, we marked MuSC nuclei with a BrdU pulse-chase strategy. α-cdKO mice received BrdU injections three times a day for five consecutive days (10-14 DPT) and TA muscles were harvested at 21 DPT then immunostained for BrdU and markers of the muscle lineage (Supplemental Fig. 6A). In control mice, 4% of Pax7^+^ cells incorporated BrdU, whereas 8% of Pax7^+^ cells in α-cdKO muscle did so, though this comparison fell short of p < 0.05 (Fig. S6B,C). It is likely that at 21 DPT, few cells that incorporated BrdU would have remained Pax7^+^. Sections were therefore assessed for expression of Myogenin, which drives terminal differentiation of muscle progenitor cells (Hernández-Hernández et al., 2017). Myogenin^+^ cells were rarely observed in TA muscle sections of control mice. In contrast, Myogenin^+^ cells were abundant in α-cdKO mice and 97% of these cells were also BrdU^+^ (Fig. 6A,B). Most (58%) BrdU^+^ cells were also positive for Myogenin (Fig. 6C). Interestingly, while almost all MyoD^+^Ki67^+^ cells in α-cdKO mice were located in the interstitial space, 33% of Myogenin^+^BrdU^+^ cells were found under the basal lamina (Figs. 5D,F and 6A,D). MyoD expression precedes and induces Myogenin expression, suggesting that these latter cells have returned to the MuSC niche.

**Fig. 6.**
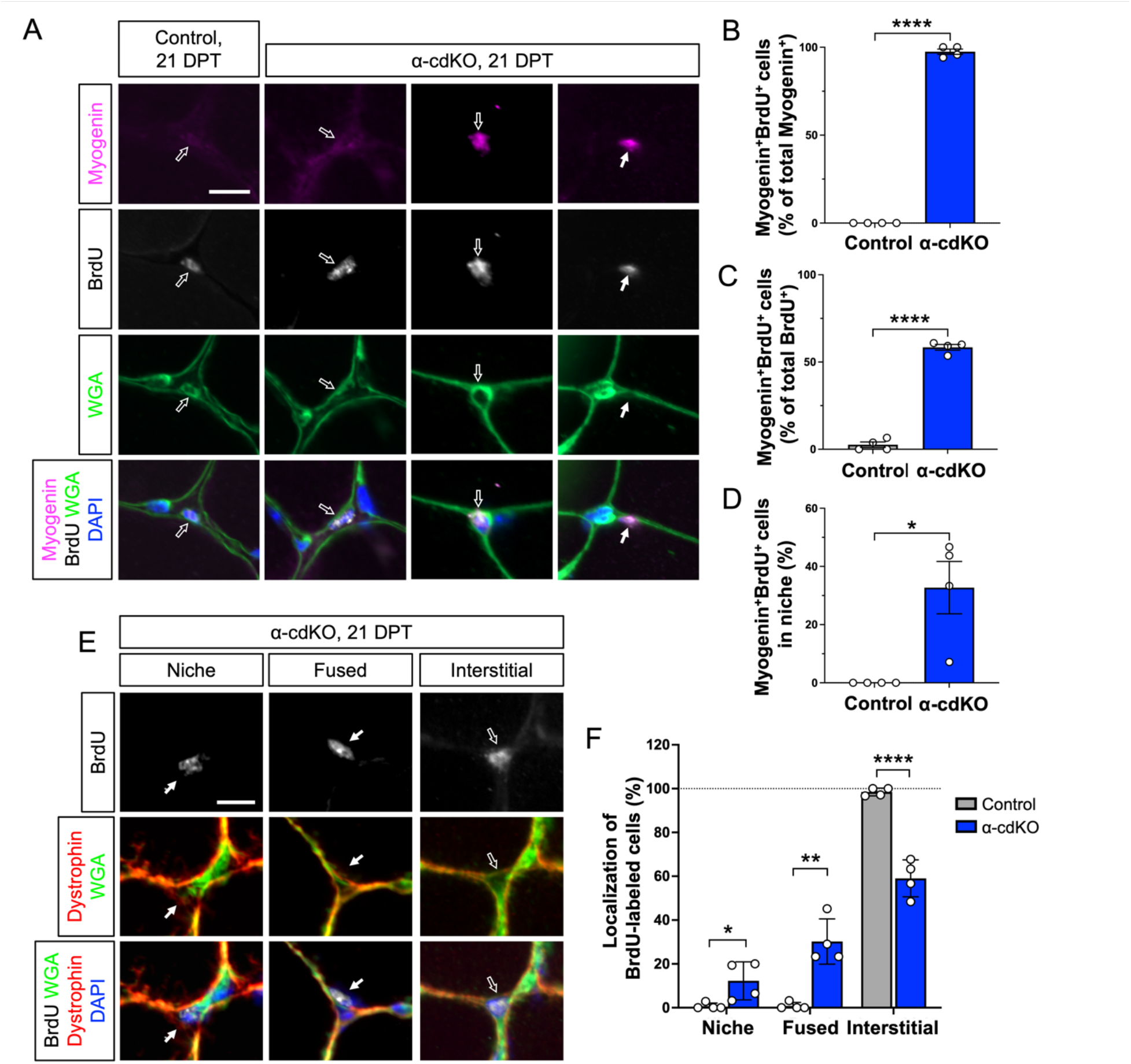
α-cdKO MuSCs are lost to precocious differentiation and fusion. **(A-D)** TA muscle sections from control and α-cdKO mice at 21 DPT were immunostained (Myogenin, magenta; BrdU, grey; WGA, green; DAPI, blue) to study cell cycle progression into S-phase by BrdU incorporation and precocious differentiation of myogenic cells by expression of Myogenin (A). Cycling myogenic cells (B,C) and location of Myogenin^+^BrdU^+^ cells (D) were quantified. **(E-F)** TA muscle sections from control and α-cdKO mice at 21 DPT were immunostained for sarcolemma marker dystrophin (red) and basal lamina marker WGA (green) (E) to assess if actively cycling myogenic cells under the basal lamina were able to initiate fusion into the myofiber (F). Scale bars: 10 μm (A,E). Each data point represents quantification of n≥30 cells (TA muscle sections) from each animal. Solid arrow denotes cell under the basal lamina; open arrow denotes interstitial cell. All data represent n=4 per genotype and represented as mean ± S.E.M with comparisons by two-tailed unpaired Student’s t-test. * = p < 0.05, ** = p < 0.01, **** = p < 0.0001.

To determine whether α-cdKO MuSCs may fuse with their associated myofibers, we stained TA muscle sections from 21 DPT mice for BrdU, plus dystrophin to mark the interior of the myofiber sarcolemma and wheat germ agglutinin (WGA) to label the basal lamina. In control mice, BrdU^+^ cells were infrequent and virtually all were located in the interstitial space (these cells are likely to be muscle-resident cells that divide at a measurable rate during homeostasis [e.g., fibroadipogenic progenitors or endothelial cells]) (Joe et al., 2010; Murphy et al., 2011; Verma et al., 2018; Wosczyna et al., 2019). In contrast, 59% of BrdU^+^ cells in α-cdKO muscle were located in the interstitial space, and as discussed above most of these expressed the muscle-specific marker Myogenin. 12% of BrdU^+^ cells were found within the typical MuSC niche, sandwiched between the basal lamina and myofiber surface, while 30% were positive nuclei found under both WGA and dystrophin, having fused with the myofiber (Fig. 6E,F). We found no evidence to suggest that cells had diluted their incorporated BrdU via proliferation. BrdU dilution to below immunofluorescent detection limits has been quantified as requiring between two to five divisions (Cutler et al., 2022; Parretta et al., 2008; Wilson et al., 2008), yet no clusters of mononucleated BrdU^+^ cells were observed. These findings are therefore consistent with the hypothesis that α-cdKO MuSCs initiate differentiation in the interstitial space, then return to the niche and fuse with the myofiber. Furthermore, this appears to be the sole mechanism of MuSC attrition in α-cdKO mice.

## DISCUSSION

A variety of adult stem cells, including MuSCs, exist in a state of quiescence for extended periods of time (Fuchs & Blau, 2020). Quiescence is an actively maintained property, requiring multiple signals provided by the stem cell niche (Urbán & Cheung, 2021; van Velthoven & Rando, 2019). Cell-cell and cell-matrix adhesion between stem cells and their niche are important regulators of this process, promoting stable niche localization as well as specific signaling information (Chen et al., 2013; Parsons et al., 2010; Polisetti et al., 2016; Schüler et al., 2022). Maintenance of MuSC quiescence is important for stem cell function; when it is broken non-physiologically (e.g., in geriatric or genetically-modified mice), it usually leads to loss of a functional MuSC pool and impaired regeneration (Evano & Tajbakhsh, 2018; Hong et al., 2022; Kimmel et al., 2020; Relaix et al., 2021). We previously reported that genetic removal from MuSCs of M-cadherin, the major niche cadherin by expression level, had no effect on MuSC homeostasis. In contrast, removal of N-cadherin resulted in a propensity of MuSCs to asynchronously enter a state of partial activation, followed by progression through full activation, cell division, cell differentiation, and fusion with myofibers (Goel et al., 2017). Combined loss of N- and M-cadherin exacerbated this phenotype (Goel et al., 2017). Despite a failure to maintain quiescence, MuSCs lacking both cadherins remained polarized and in a normal niche location, and they preserved regenerative and self-renewal capabilities. We attributed these unusual latter properties to yet additional cadherins expressed at lower levels than N- and M-cadherin (Yue et al., 2020). Nevertheless, these observations raised the possibility that while cadherins are clearly important for maintenance of MuSC quiescence, they may be dispensable for stable niche localization. Basal adhesion alone may be sufficient, and alternative apical adhesion mechanisms may also exist, including Notch ligand/receptor interactions and yet-to-be-explored factors such as protocadherins (Murata et al., 2014; Pancho et al., 2020). We therefore addressed the effects in MuSCs of complete loss of cadherin-based adhesion by genetically removing pairs of redundant catenin proteins that are required for cadherin function. Our findings demonstrate that cadherin-based adhesion is essential for stable niche localization of MuSCs, and that MuSCs are depleted in its absence. Furthermore, N-cadherin plays a specific role in preventing MuSCs from breaking quiescence, a role that is distinct from the requirement for cadherins in maintaining niche adhesion.

MuSC-specific removal of β- and γ-catenins or, separately, αE- and αT-catenins led to MuSC attrition over the course of 4 weeks in both lines. A fraction of MuSCs persisted at this and later timepoints but the majority of such cells were still positive for at least one of the target proteins, indicating incomplete Cre-mediated recombination or exceptionally long perdurance of the protein in a subset of MuSCs. While these catenin pairs are essential for cadherin-dependent cell-cell adhesion, they also play additional roles in cell regulation. β-catenin is a critical signal transducer of canonical Wnt signaling, and γ-catenin plays a central role in organization of desmosomes via binding to desmosomal cadherins (Green et al., 2019; van der Wal & van Amerongen, 2020). However, canonical Wnt signaling is silent during MuSC quiescence, and MuSCs do not express desmosomal cadherins (Murphy et al., 2014; Parisi et al., 2015; Rudolf et al., 2016; Yue et al., 2020). Additionally, single genetic removal of β- or γ-catenin was without effect on MuSC numbers at homeostasis, whereas combined removal resulted in MuSC attrition (Murphy et al., 2014; Rudolf et al., 2016). These results indicate that β- and γ-catenin share a function required for MuSC maintenance, and that this function is almost certainly classical cadherin-based adhesion. This conclusion is consistent with results from ablation of these two cadherin-binding proteins in heart muscle and in motor neuron pools during spinal cord development (Demireva et al., 2011; Swope et al., 2012). α-catenins also play roles in regulating signaling, but these are usually performed as components of adhesion complexes (Priya & Yap, 2015; Yap et al., 2018). For example, in some cell types α-catenin binds YAP to prevent its dephosphorylation and nuclear translocation (Schlegelmilch et al., 2011; Silvis et al., 2011). However, YAP expression is induced later during MuSC activation, and expression of constitutively active YAP in adult MuSCs in vivo is insufficient to break quiescence (Judson et al., 2012; Tremblay et al., 2014). Release of YAP as a consequence of adhesion complex deficiency is therefore unlikely to explain the phenotype of α-cdKO mice. We therefore conclude that the primary defect in these mouse lines is a loss of cadherin-based adhesion in MuSCs and not other functions of catenins.

It is intriguing that MuSCs lacking N- and M-cadherin and α-cdKO MuSCs both appear to go through similar stages of activation and progression in the absence of injury, including expressing MyoD, entering the cell cycle, differentiating, and fusing with myofibers. However, N- /M-cadherin-deficient MuSCs remain in the niche and are not depleted, whereas α-cdKO MuSCs exit the niche and are depleted (Fig. 7). It may be that the ability of N-/M-cadherin mutant MuSCs to stay under the basal lamina, due to a diminished but functional complement of classical cadherins, allows them to enter and slowly complete a normal activation process. In contrast, full loss of cadherin-based adhesion in α-cdKO MuSCs results in escape from the niche, where the interstitial microenvironment may lack signals required for normal activation and myogenic progression, ultimately leading to precocious differentiation and MuSC attrition. Cadherin-dependent niche localization is not, however, required for MuSC survival or maintenance of myogenic identity, as the sole fate of α-cdKO MuSCs appears to be differentiation and fusion with myofibers. The latter event requires these cells to recross the basal lamina. It is possible, in fact, that α-cdKO MuSCs cross the basal lamina multiple times prior to fusion. This may be related to expression of matrix metalloproteinases by activated MuSCs (Pallafacchina et al., 2010), but other mutants display similar unscheduled activation without exiting the niche (Bjornson et al., 2012; Mourikis et al., 2012; Rozo et al., 2016).

**Fig. 7.**
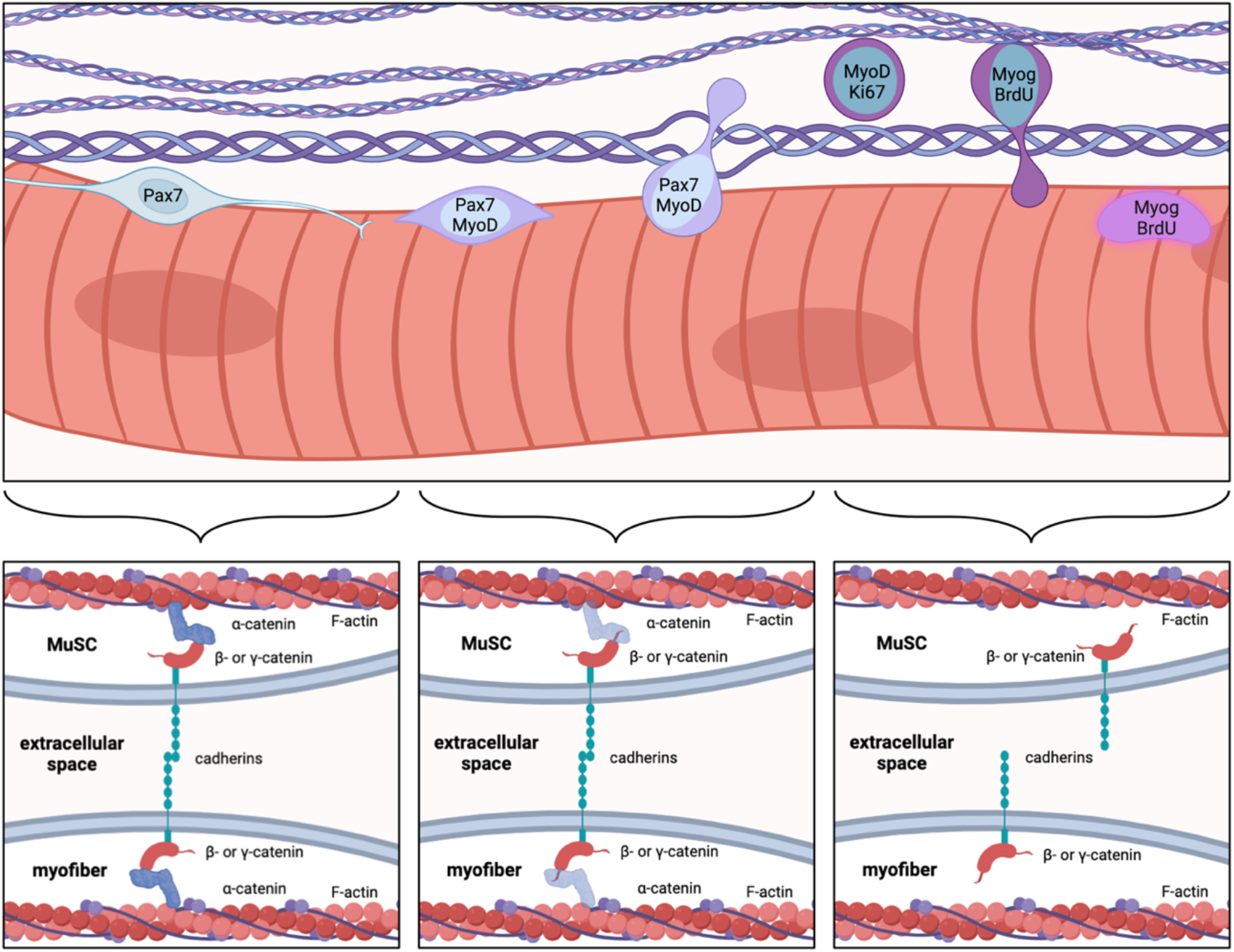
Schematic representation of α-cdKO MuSC behavior. Attrition of Pax7^+^ MuSCs in α-cdKO mice is due to loss of apical, cadherin-catenin–based adhesion to the myofiber. This results in cells that activate in the absence of injury, exit the niche, enter the cell cycle, and terminally differentiate to fuse into the myofiber. Made with BioRender.

These observations raise fundamental questions: what are the adhesion requirements for MuSCs to adopt stable niche localization, and are they the same as those necessary to maintain quiescence? The basal lamina is mainly composed of laminin and collagen IV (Baghdadi et al., 2018; Schüler et al., 2022). β1-integrin heterodimerizes with α7-integrin to form the major MuSC laminin receptor and is also a component of collagen receptors (Rozo et al., 2016; Schüler et al., 2022; Mashinchian et al., 2018). MuSC-specific mutation of β1-integrin results in loss of cell polarity and MuSC attrition but the cells remain in the niche, under the basal lamina. Although potential mechanisms for basal adhesion that do not rely on β1-integrin may exist in MuSCs, their roles are not clear (Mashinchian et al., 2018). Apical adhesion mechanisms may therefore be sufficient to provide niche localization but are insufficient to maintain cell polarity and quiescence in the absence of β1-integrin-containing adhesion receptors.

Our findings with catenin-deficient MuSCs indicate that cadherin-based adhesion is required for both niche localization and maintenance of quiescence. Exit from the niche is not, however, an automatic consequence of MuSC activation in the absence of injury. In addition to the example of β1-integrin, conditional mutation of the Notch-responsive transcriptional regulator RBP-J leads to MuSC activation and attrition via precocious differentiation, while the cells remain in the niche (Bjornson et al., 2012; Mourikis et al., 2012). Additionally, the Notch target collagen V signals via calcitonin receptor (CalcR) in MuSCs (Baghdadi et al., 2018). This interaction is required for MuSC quiescence but only 4% of CalcR-deficient MuSCs are present outside the basal lamina, about 10-fold less than seen with α-cdKO MuSCs (Yamaguchi et al., 2015). Therefore, the interstitial localization of α-catenin-deficient MuSCs is likely due directly to loss of cadherin-based adhesion to the adjacent myofiber, rather than a secondary consequence of MuSC activation.

Taken together, our current and previous findings (Goel et al., 2017; Kann et al., 2022) demonstrate that cadherin-based adhesion is critical for niche localization and preservation of MuSCs. N-cadherin is specifically required for maintenance of MuSC quiescence independent of niche localization. Quiescent MuSCs have long, heterogeneous projections that appear to serve as sensors of the stem cell niche, as they rapidly retract upon muscle injury (Kann et al., 2022; Ma et al., 2022). N-cadherin protein is enriched on projections, and N-cadherin-deficient MuSCs lack or have significantly shorter projections (Kann et al., 2022). As N-cadherin is dispensable for niche localization, we hypothesize that its major role in regulating MuSC quiescence is promoting outgrowth and/or maintenance of projections, with a minimum homeostatic projection length required for quiescence. Anchorage to the niche must be provided either by different, or the full complement of, cadherins, as shown here by the phenotype of MuSCs lacking catenins.

Cadherin-catenin complexes are best understood as factors bridging cell-cell contact and adhesion with cytoskeletal structure and dynamic regulation of cell tension (Leckband & de Rooij, 2014; Mège & Ishiyama, 2017). Work on MuSC cellular projections suggests that the first step in injury-induced MuSC activation may be a change to the biomechanical environment of the niche (Krauss & Kann, 2023). Additionally, changes to the actin and microtubule cytoskeletons are among the earliest events visible after MuSC activation (Kann et al., 2022). How these cytoskeletal structures differ between quiescent MuSCs and MuSCs activated by injury or by partial or complete loss of cadherin-catenin–dependent adhesion is now important to address. This will require new preparations and high-resolution tools, as isolated MuSCs do not retain a physiological morphology and myofiber sarcomeres interfere with clear visualization of MuSC actin structures on single myofiber preparations (Kann et al., 2022). Our findings indicate that various types of cadherin-catenin–based adhesion are required in MuSCs for both niche localization and quiescence, and can be distinguished from the role of integrin-based adhesion, suggesting that such approaches will provide significant insight into stem cell niche function.

## MATERIALS AND METHODS

### EXPERIMENTAL MODEL AND SUBJECT DETAILS

#### Animals

Mice were housed and maintained in accordance with recommendations set in the Guide for the Care and Use of Laboratory Animals of the National Institutes of Health. All animal protocols used in these studies were approved by the Icahn School of Medicine at Mount Sinai Institutional Animal Care and Use Committee (IACUC).

#### Mouse breeding

To generate the *Ctnna1^flox/flox^*;*Ctnna3^flox/flox^*;*Pax7 ^tm1(CreERT2)Gaka/+^* line, 8-12 week old homozygous *Ctnna1^flox/flox^*;*Ctnna3^flox/flox^*mice (Li et al., 2015, 2012; Vasioukhin et al., 2001) were crossed with 8-12 week *Pax7^tm1(CreERT2)Gaka^*heterozygous mice (Murphy et al., 2011) (Jackson Laboratory, 017763). The offspring were crossed until litters yielded mice homozygous for floxed alleles of αE- and αT-catenin, and heterozygous for *Pax7 ^tm1(CreERT2)Gaka^*. These mice were designated as α-cdKO. Littermates lacking *Pax7 ^tm1(CreERT2)Gaka^* were used as controls. To estimate the recombination efficiency of the system, we made the assumption that any MuSCs that were no longer detectable via Pax7 staining had undergone full recombination, i.e., lacking expression of all four α-catenin alleles and degradation of perduring protein via homeostatic turnover. Minimum recombination efficiency by this definition is 78%, the percentage of MuSCs lost in double-mutant mice compared to controls. Of the 22% of MuSCs remaining on fibers, 82% were αE-catenin-immunoreactive and 78% were αT-catenin-immunoreactive. We therefore conclude that 18% of the remaining 22% of MuSCs (4% of the starting population) have a complete loss of catenin protein. This yielded an estimated 82% total recombination efficiency in α-cdKO mice at 28 DPT.

To generate the *Ctnna1^flox/flox^*;*Ctnna3^flox/flox^*;*Pax7 ^tm1(CreERT2)Gaka/+^*;*ROSA26^LSL-tdTomato^* line, α-cdKO mice were crossed with 8-12 week old homozygous *Gt(ROSA)26Sor^tm14(CAG-tdTomato)Hze^* mice purchased from Jackson Laboratory (Madisen et al., 2010) (Jackson Laboratory, 007914).

To generate the *Ctnnb1 ^tm2Kem/tm2Kem^*;*Jup ^tm1.1Glr^*^/*tm1.1Glr*^;*Pax7 ^tm1(CreERT2)Gaka/+^* line, 8-12 week old homozygous *Ctnnb1^tm2Kem^*;*Jup^tm1.1Glr^*mice provided by Glenn Radice (Li et al., 2011; Swope et al., 2012) (Jackson Laboratory, 004152 and 017575) were crossed with 8-12 week *Pax7^tm1(CreERT2)Gaka^* heterozygous mice (Murphy et al., 2011) (Jackson Laboratory, 017763). The offspring were crossed until litters yielded mice homozygous for floxed alleles of β- and γ-catenin, and heterozygous for *Pax7 ^tm1(CreERT2)Gaka^*. These mice were designated as βγ-cdKO. Littermates lacking *Pax7 ^tm1(CreERT2)Gaka^* were used as controls. Estimation of recombination frequency was calculated as above for α-cdKO mice. The percentage of MuSCs lost in double-mutant mice was 71% and of the remaining 29%, 53% were β-catenin-immunoreactive and 66% were γ-catenin-immunoreactive. Therefore, 34% of the remaining 29% of MuSCs (10% of the starting population) have lost β- and γ-catenin, yielding an estimated 81% total recombination efficiency in βγ-cdKO mice at 28dpt.

To generate the *Jup^tm1.1Glr^*;*Pax7 ^tm1(CreERT2)Gaka/+^* line, 8-12 week old *Ctnnb1^tm2Kem/tm2Kem^*;*Jup^tm1.1Glr^*^/*tm1.1Glr*^;*Pax7 ^tm1(CreERT2)Gaka/+^* mice were crossed with 8-12 week *Pax7^tm1(CreERT2)Gaka/+^*mice (Murphy et al., 2011) (Jackson Laboratory, 017763) until resultant litters had homozygous wild type alleles for β-catenin, homozygous floxed alleles of γ-catenin, and heterozygous for *Pax7 ^tm1(CreERT2)Gaka^.* These mice were designated as γ-cKO. Littermates lacking *Pax7 ^tm1(CreERT2)Gaka^*were used as controls. *Pax7 ^tm1(CreERT2)Gaka^; ROSA26^LSL-tdTomato^*mice were utilized in FACS and scRNAseq experiments as controls and were denoted ‘Control’ in single cell RNA sequencing experiments.

Primers used for PCR genotyping are listed in Table S1.

### METHOD DETAILS

#### Tamoxifen administration

Adult mice (2-3 months) were injected intraperitoneally for 5 consecutive days with 125 mg/kg of tamoxifen dissolved in corn oil (Toronto Research Chemicals, T006000; ThermoFisher Scientific, 405435000). Tamoxifen supplementation in chow was used during injury experiments as noted in the main text and figure legends(Envigo TD,130857).

#### Muscle isolation

Whole TA muscles and single myofibers from EDL muscles were isolated as previously described (Goel & Krauss, 2019). Briefly, mice were sacrificed via CO_2_ inhalation and death was confirmed with cervical dislocation. Skin was removed from the hindlimbs from the ankle joint to the hip. Whole TA muscles were mounted in 10% w/v tragacanth gum (Alfa Aesar, A18502) and snap frozen in 2-methylbutane (Fisher Scientific, O3551-4) before being transferred to a -80°C freezer for storage. Frozen TA muscles were cryosectioned at 10μm thickness and collected on positively charged slides. Sections were kept at -20°C until used for immunostaining or histology.

The tendons of the EDL were cut at the ankle and knee, and the EDL was immediately incubated in plating medium (DMEM + 10% horse serum + 2% Penicillin/Streptomycin + 1% HEPES) containing type I collagenase (2.6 mg/mL; Gibco, 17100-017) in a 37°C shaking water bath for 53 minutes, followed by trituration with a wide bore glass pipet in plating medium in a fresh 10 cm plate. Fibers were allowed to rest in a 37°C incubator for 5 minutes before they were collected for fixation in a final concentration of 4% paraformaldehyde in PBS for 10 minutes. Myofibers were washed with fresh PBS after fixation and maintained at 4°C until used for immunostaining.

#### Skeletal Muscle injury

An acute barium chloride (BaCl_2_) injury model was adapted and used to study muscle regeneration (Tierney & Sacco, 2016). Fur was removed from the injection site of the anterior hindlimb with clippers and disinfected with ethanol. A 1 mL syringe equipped with a 30g needle was used to pierce the skin nearly parallel to the ankle and inserted through the length of the TA, stopping just short of the knee. A repetitive process of slowly releasing 1.2% BaCl_2_ dissolved in normal saline (Thermo Scientific, 612281000) and withdrawing the needle tip slightly ensured adequate diffusion of BaCl_2_ into the muscle and minimized leakage. A total of 75 μL was injected along the length of the TA for full muscle injury. Mice were allowed to recover fully from anesthesia on a heating pad. Four weeks after the initial injury, mice were re-injured, and muscles were harvested at 7 and 28dpi.

#### BrdU pulse-chase labeling

Mice were injected intraperitoneally three times daily for five consecutive days with 5-bromo-2’-deoxyuridine (BrdU, Sigma B9285-1G) diluted to 30 mg/kg in PBS. Mice were harvested after a chase period of 1 week to label cells undergoing S phase entry *in vivo*.

#### Immunofluorescence on single myofibers

After single EDL myofibers were acquired as described above, fixed myofibers were permeabilized with 0.2% TritonX-100 in PBS (PBST) for 10 minutes, followed by incubation with 10% goat serum in staining media for 1 hour on a rotating shaker at room temperature. Primary antibodies were added and fibers were incubated overnight at 4°C on a shaker. Fibers were washed with PBS and PBST before blocking in 10% goat serum. Secondary antibodies were added to incubate for an hour in the dark at room temperature before being washed again with PBS and PBST and mounted on positively charged slides with DAPI Fluoroshield mounting media (Abcam ab104139). Antibodies and dilutions used for immunofluorescent staining can be found in Table S2.

#### Immunofluorescence on whole muscle cross sections

Frozen TA sections were allowed to equilibrate to room temperature and fixed in 4% paraformaldehyde in the dark for 20 minutes. The slides were rinsed in PBS, permeabilized in -20°C methanol for 6 minutes, and washed with PBS. Antigen retrieval was performed using 0.01M citric acid buffer, pH 6.0, at 90°C for 10 minutes. Slides were washed again in PBS, blocked in 5% BSA in PBS for 2 hours, and primary antibodies were added to incubate overnight at 4°C. Slides were washed in PBS and blocked with 0.1% BSA in PBS for 30 minutes. Secondary antibodies were added and slides incubated for 1 hour at room temperature, washed with PBS, and mounted with DAPI Fluoroshield mounting media (Abcam, ab104139).

#### TUNEL Assay

The Click-iT™ Plus TUNEL Assay Kit for In Situ Apoptosis Detection 594 (ThermoFisher, C10618) was used to detect DNA breaks on frozen TA sections. The manufacturer protocol was used. Sections were fixed in 4% PFA for 15 minutes at 37°C, then washed in 1X PBS twice for 5 minutes. Sections were then covered and incubated with Proteinase K solution for 15 minutes at room temperature before washing with PBS once for 5 minutes. Sections were fixed a second time in 4% PFA for 5 minutes at 37°C, followed by two washes of PBS for 5 minutes and a rinse in deionized, RNase-free water. A positive control was prepared by inducing DNA breaks with DNaseI solution for 30 minutes at room temperature before a deionized water rinse. All sections were then incubated in TdT reaction buffer for 10 minutes at 37°C, followed by the TdT reaction mixture at 37°C for an hour. Sections were washed with 3% BSA and 0.1% Triton X-100 in PBS for 5 minutes at room temperature and rinsed with PBS after. Sections were then treated with Click-iT™ Plus TUNEL reaction cocktail for 30 minutes at 37°C in the dark, washed with 3% BSA in PBS for 5 minutes, and rinsed with PBS. The TUNEL reaction was followed by antibody staining. Sections were blocked for an hour at room temperature in 3% BSA in PBS and incubated with an anti-laminin antibody overnight at 4°C in the dark. Sections were washed with 3% BSA in PBS twice for five minutes and incubated with Alexa Fluor 488 anti-rabbit IgG secondary antibody for 30 minutes at room temperature. Sections were washed with 3% BSA in PBS twice for five minutes, dried, and mounted with DAPI Fluoroshield mounting media (Abcam, ab104139).

#### Picrosirius Red staining

Frozen TA sections were air dried overnight at room temperature. Before staining, sections were circled with a Pap pen (Vector Laboratories, H-4000) and allowed to dry. A hybridization chamber with a water reservoir was used as a humidity chamber and set to 56°C. The slides were added to the chamber and a disposable pipette was used to drop Bouin’s fixative onto the slides and incubated for 15 minutes. Bouin’s fixative was removed in a fume hood and slides were washed in distilled water twice before placing them into a coplin jar with 30 mL of 0.1% Sirius Red in saturated picric acid (Electron Microscopy Services, 26357-02) and incubating for 2 hours on a shaker. Slides were washed in 0.5% acetic acid, dehydrated in ethanol, equilibrated in xylenes for 10 minutes, and mounted with Cytoseal XYL (Fisher Scientific, 22-050-262).

#### FACS isolation of MuSCs

Single cell suspensions from mouse hindlimb whole muscle were acquired with modifications from a protocol described previously (Gromova et al., 2015). After euthanasia, skin was removed from both hindlimbs, and whole hindlimbs were removed to the hip and placed on ice in PBS. Muscles were excised with care to remove fat, tendons, blood vessels, nerves, and bones. Muscles were minced with a sterile razor blade and transferred into digestion medium containing collagenase II and dispase II (Fisher Scientific, 17-101-015; 17-105-041) for enzymatic dissociation of mononuclear cells from muscle fibers in a rotating water bath at 37°C. A 10 mL syringe with a 20 g needle was used further release mononuclear cells into suspension before being passed through 40 μm cell strainers. Cells were pelleted at 300 g for 5 minutes before washing with ice cold Hanks’ Balanced Salt Solution without magnesium and calcium supplemented with 2% FBS and 1 mM EDTA (HBSS+), pelleting again, and resuspending in 4ml of HBSS+. Samples were kept on ice and counterstained with SYTOX Blue before FACS isolation of MuSCs. Each sample was pooled from two littermate mice of the same genotype with 2 samples sequenced per genotype. A Sony MA900 equipped with a 100 μm nozzle and 20 psi pressure was used for FACS isolation.

#### Single cell RNA sequencing analysis processing

Sequenced fastq files were aligned, filtered, barcoded and UMI counted using Cell Ranger Chromium Single Cell RNA-seq version 7.1.0, by 10X Genomics with Cell Ranger, GRCm38 database (version 2020-A) as the mouse genome reference. Each dataset was filtered to retain cells with ≥ 1000 UMIs, ≥ 400 genes expressed, and <15% of the reads mapping to the mitochondrial genome. UMI counts were then normalized so that each cell had a total of 10,000 UMIs across all genes and these normalized counts were log-transformed with a pseudocount of 1 using the “LogNormalize” function in the Seurat package. The top 2000 most highly variable genes were identified using the “vst” selection method of “FindVariableFeatures” function and counts were scaled using the “ScaleData” function. Datasets were processed using the Seurat package (version 4.0.3) (Hao et al., 2021).

Principal component analysis was performed using the top 2000 highly variable features (“RunPCA” function) and the top 30 principal components were used in the downstream analysis. Datasets for each patient were integrated by using the “RunHarmony” function in the harmony package (version 0.1.0) (Korsunsky et al., 2019). K-Nearest Neighbor graphs were obtained by using the “FindNeighbors” function whereas the UMAPs were obtained by the “RunUMAP” function. The Louvain algorithm was used to cluster cells based on expression similarity at 0.2 resolution.

Differential markers for each cluster were identified using the Wilcox test (“FindAllMarkers” function) with adjusted p-value < 0.05 and absolute log2 fold change > 0.25, and 1,000 cells per cluster were randomly picked to represent each cluster. Top two significant markers in terms of log2 fold change for all clusters were plotted using Seurat DotPlot command. The top up-regulated genes and curated genes from the literature were used to assign cell types to the clusters. Unicell Deconconvolve was used to annotate the cells using the log normalized counts of the integrated dataset (version 0.0.3) (Charytonowicz et al., 2023). Prediction values for each cell type are merged with the original Seurat object. UMAPs of prediction value for skeletal and striated muscle are generated using the Seurat Featureplot function (version 4.0.3) (Hao et al., 2021).

P values for cell proportions within clusters were calculated using a logistic regression method. The lme4 package was used to fit data to a logistic regression model (version 1.1-27) (Bates et. al 2015). The odds ratio and p-value of seeing this ratio between conditions was calculated using the emmeans package(version 1.6.2) (Lenth 2021).

### QUANTIFICATION AND STATISTICAL ANALYSIS

#### Single myofiber imaging and quantification

Images were collected using 100x/1.40 oil objectives with up to an additional 2x digital zoom on a Leica DMI SP8 inverted confocal microscope equipped with Leica Application Suite. Line averaging was used to improve signal-to-noise ratio in representative images as follows: line average-3, frame average-2. Images were exported using the native Leica Application Suite as TIFFs and imported to ImageJ for post-imaging analysis with adjustment of brightness and contrast only. For quantification of cell numbers and presence of catenin protein, as indicated by immunofluorescent signal, at least 10 fibers were analyzed per mouse with an n≥4 per genotype, as noted in figure legends.

#### Whole muscle cross section imaging and quantification

Widefield images were collected using a 20x/0.75 air objective on a Zeiss AxioImager Z2 equipped with Zen Blue Software. Images were exported using the native Zeiss Application Suite as .czi files or TIFFs and imported to ImageJ for post-imaging analysis with adjustment of brightness and contrast only.

For quantification of cell numbers during homeostasis and regeneration, 10 random fields of view were taken at 20x magnification per mouse with an n≥3 per genotype, as noted in figure legends. For quantification of cell localization, cell expression of activation markers, and BrdU incorporation, a minimum of 30 cells were counted per animal with an n≥3 per genotype, as noted in figure legends. Picrosirius Red staining was quantified using ImageJ where images were segmented by red-stained collagen and measuring percent coverage by the segmented area. Myofiber minimum feret diameter was quantified using the ImageJ package MuscleJ (Mayeuf-Louchart et al., 2018).

#### Statistical analyses

All experiments, excluding FACS isolation and single cell RNA-sequencing experiments, were performed using mice with n≥3 per genotype as specified in figure legends. Mean, standard error of means (SEM), 95% confidence intervals, unpaired two-tailed Student’s t-tests, and Mann-Whitney tests were calculated with Prism GraphPad Software. All graphs were generated with Prism GraphPad Software or R. Supplemental Figure 1 shares control and βγ-cdKO data from Figure 1.

## ACKNOWLEDGEMENTS

We thank Drs. Philippe Soriano and Allison Kann for their helpful comments on the manuscript and throughout the course of this work. We gratefully acknowledge Drs. Frans van Roy and Jolanda van Hengel for providing the *Ctnna3* mutant mice. We also thank the staff members of the Columbia Stem Cell Initiative Flow Cytometry Core Facility under the leadership of Michael Kissner at Columbia University Irving Medical Center, the Microscopy and Advanced Imaging CoRE at the Icahn School of Medicine at Mount Sinai, and the Genomics Core Facility at the Icahn School of Medicine at Mount Sinai for their contributions to the work presented in this manuscript.

## AUTHOR CONTRIBUTIONS

Conceptualization, M.H., A.G.B, G.R, R.S.K.; methodology, M.H., R.S.K.; validation and formal analysis, M.H., H.F.L., D.D., G.D.; investigation, M.H., H.F.L.; resources, A.G.B., G.R., D.H.; writing – original draft preparation, M.H., R.S.K.; writing – review and editing M.H., H.F.L., A.G.B., D.D., G.D., D.H., G.R., R.S.K.; visualization, M.H., D.D., G.D.; supervision, D.H., R.S.K.; funding acquisition, M.H., R.S.K. All authors approved the final manuscript.

## RESOURCE AVAILABILITY

### Lead Contact

Further information and requests for resources and reagents should be directed to and will be fulfilled by the lead contact, Robert S. Krauss.

### Materials Availability

This study did not generate new unique reagents.

### Data and code availability

Data reported in this paper has been deposited with accession number xxxxx.

This paper does not report original code.

## Supplemental Figure Legends

**Fig. S1.**
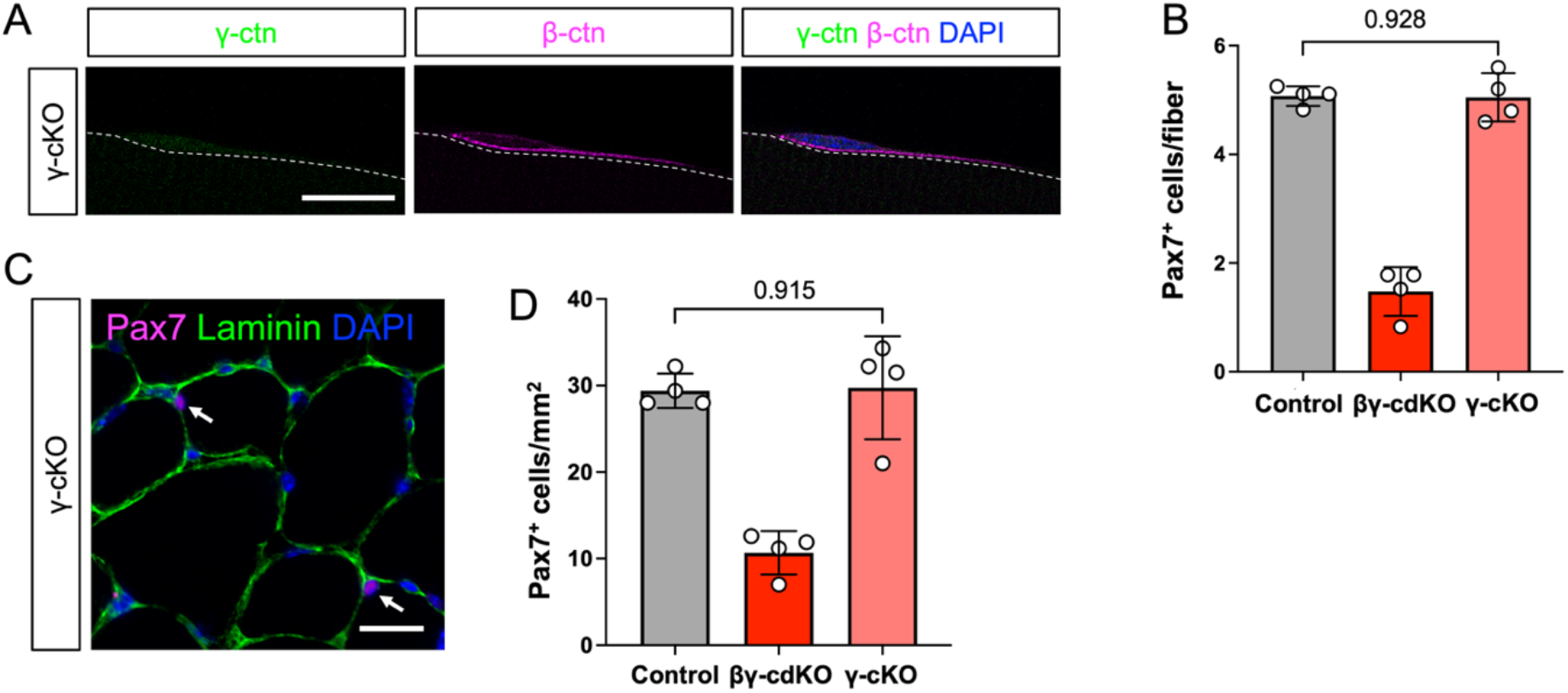
Genetic removal of γ-catenin does not lead to MuSC attrition or alter β-catenin localization. **(A-B)** Immunostaining of single myofibers from 28 DPT γ-cKO mice (γ-catenin, green;β-catenin, magenta, DAPI, blue) was used to assess loss of γ- and/or β-catenin protein (A) and MuSC attrition (B). Scale bar: 10 μm. **(C-D)** TA muscle sections from 28 DPT γ-cKO mice were immunostained (Pax7, magenta; laminin, green; DAPI, blue) and Pax7-expressing MuSCs were labeled (C) and quantified (D). Scale bar: 25 μm. Each data point represents the average from at least ten myofibers (EDL single myofibers) or ten fields (TA muscle sections) from each animal. Data represent n=4 per genotype per timepoint and represented as mean ± S.E.M with comparisons by two-tailed unpaired Student’s t-test.

**Fig. S2.**
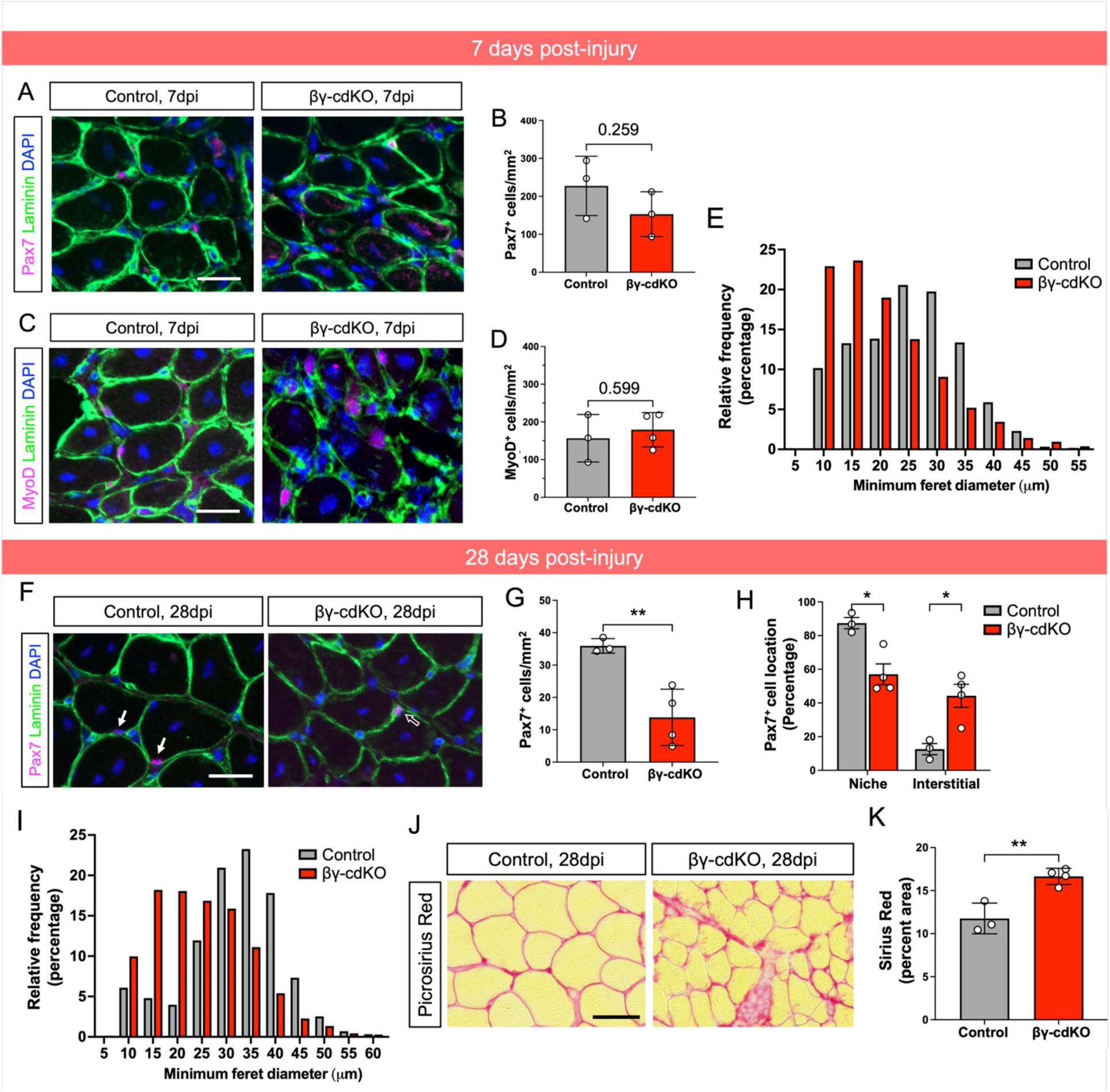
α-cdKO mice have modest defects in muscle regeneration. **(A-B)** Sections of TA muscle from control and α-cdKO mice at 28dpi with Picrosirius Red (A) to quantify collagen deposition after injury (B). Scale bar: 25 μm. **(C-D)** Average number of myofibers per field of view at 20x magnification was quantified (C), as was the average fiber diameter (D) from control and α-cdKO mice at 28dpi. Both correlated with the significant decrease in average myofiber size in Figure 2J,K. Each data point represents the average from ten fields (TA muscle sections) from each animal. Data represent n=4 per genotype per timepoint and represented as mean ± S.E.M with comparisons by two-tailed unpaired Student’s t-test.

**Fig. S3.**
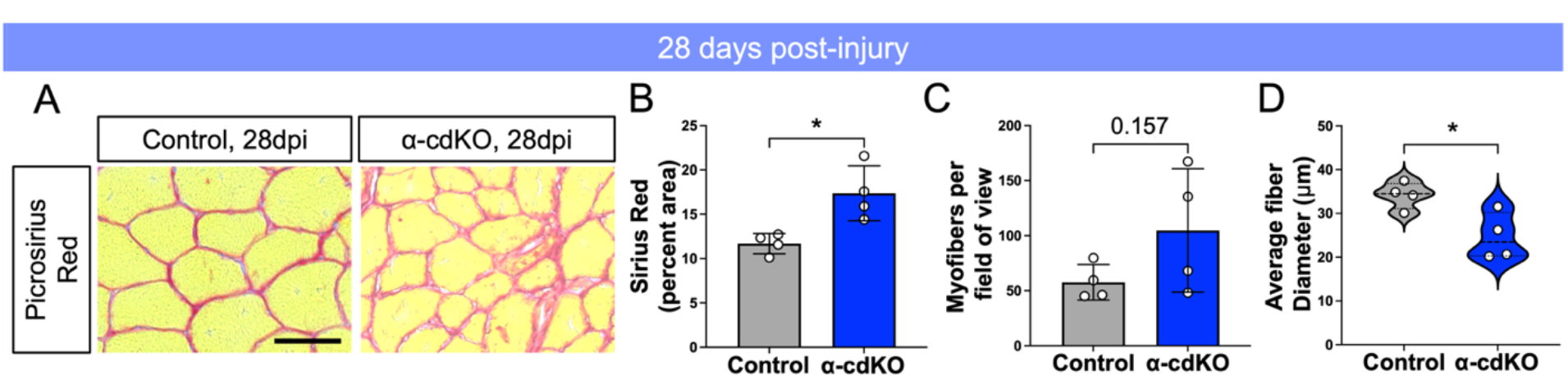
βγ-cdKO mice have modest defects in muscle regeneration. **(A-E)** At 7dpi, TA muscle sections from control and βγ-cdKO mice were immunostained (Pax7 or MyoD, magenta; laminin, green; DAPI, blue) for Pax7^+^ (A,B) and MyoD^+^ (C,D) cells to assess MuSC and myogenic progenitor numbers during early regeneration. Myofiber size was quantified by minimum feret diameter in (E). **(F-J)** At 28dpi, TA muscle sections from control and βγ-cdKO mice were immunostained (Pax7, magenta; laminin, green; DAPI, blue) (F) to assess MuSC numbers (G) and location (H) after complete regeneration. Myofiber size was quantified by minimum feret diameter in (I). Sections of TA muscle from control and βγ-cdKO mice at 28dpi with Picrosirius Red (J) to quantify collagen deposition after injury (K). Scale bars: 25 μm (A,C,F,I). Solid arrow denotes cell under the basal lamina; open arrow denotes interstitial cell. Each data point represents the average from ten fields from each animal. All data represent n≥3 per genotype and represented as mean ± S.E.M with comparisons by two-tailed unpaired Student’s t-test. ** = p < 0.01

**Fig. S4.**
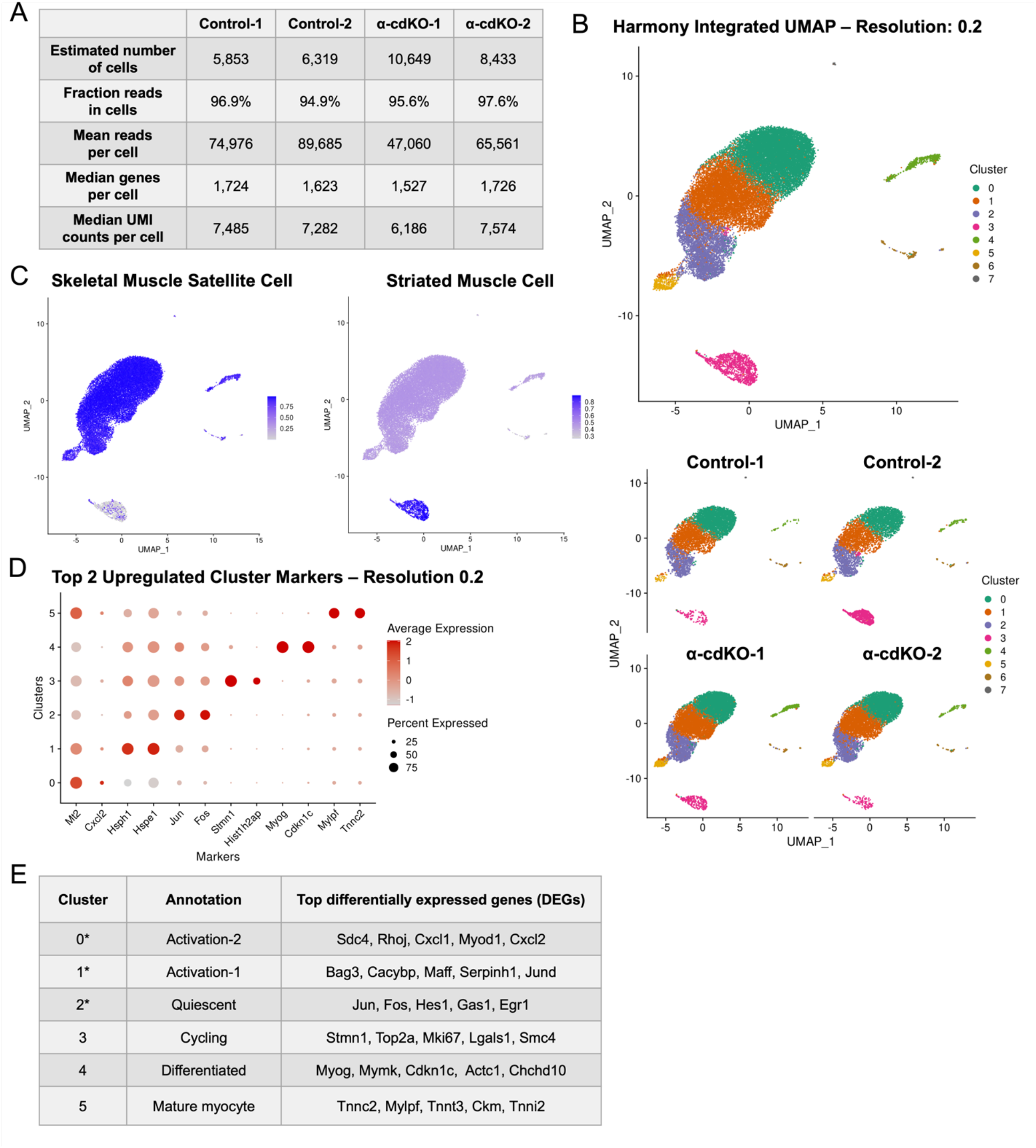
Quality control during cell purification and genomic analyses. **(A)** Single cell RNA sequencing quality control of tdTomato^+^ cells from control and α-cdKO at 14 DPT was assessed using CellRanger. **(B)** tdTomato^+^ cells from control and α-cdKO at 14 DPT were integrated using Harmony and unsupervised clustering yielded a UMAP with eight cell clusters, none of which were unique to one genotype. **(C)** UniCell Deconvolve cell type predictions score for MuSC (left) and striated muscle cell types (right). A much smaller proportion of cells were classified as mature myocytes (‘Striated Muscle Cell’) compared to MuSCs (‘Skeletal Muscle Satellite Cell’). **(D-E)** Differentially upregulated genes per cluster (D) indicate isolation-induced stress response in some clusters, but used in conjunction with other differentially expressed genes per cluster, allowed author annotation of cluster identities (E).

**Fig. S5.**
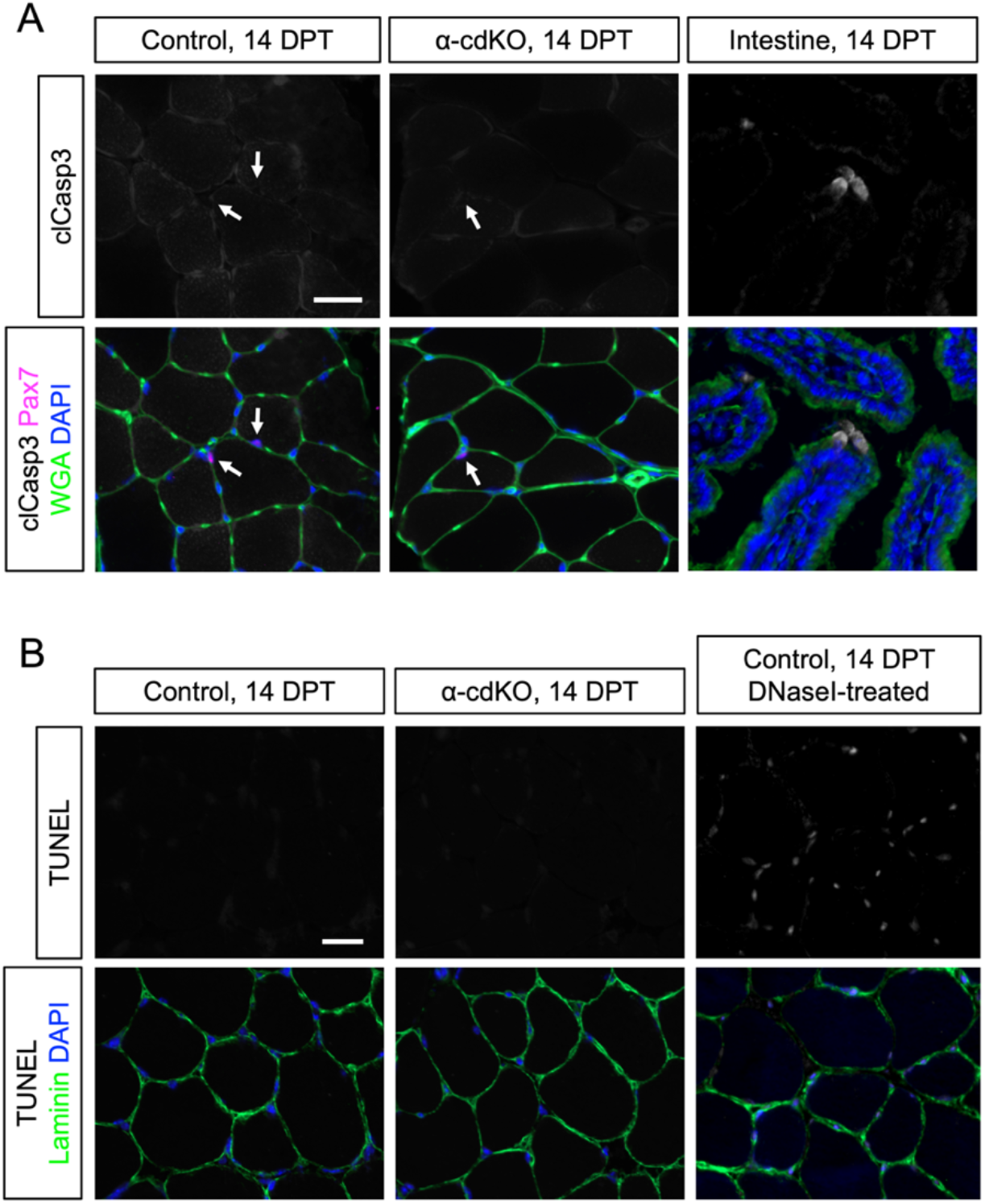
Lack of evidence for apoptosis of α-cdKO MuSCs. **(A)** Immunostaining of sections through TA muscle of control and α-cdKO mice at 14 DPT for Pax7 (magenta), cleaved Caspase-3 (grey), WGA (green), and DAPI (blue). Intestine used as a positive control. Arrows denote location of Pax7^+^ cells. No evidence of cleaved Caspase-3 was found in muscle sections was found. Scale bar: 25μm. **(B)** TUNEL detection (grey) with immunostaining of sections through TA muscle of control and α-cdKO mice at 14 DPT for Laminin (green), and DAPI (blue). Control TA muscle section treated with DNaseI used as a positive control. No evidence of TUNEL detection of DNA breakage was found. Scale bar: 25μm.

**Fig. S6.**
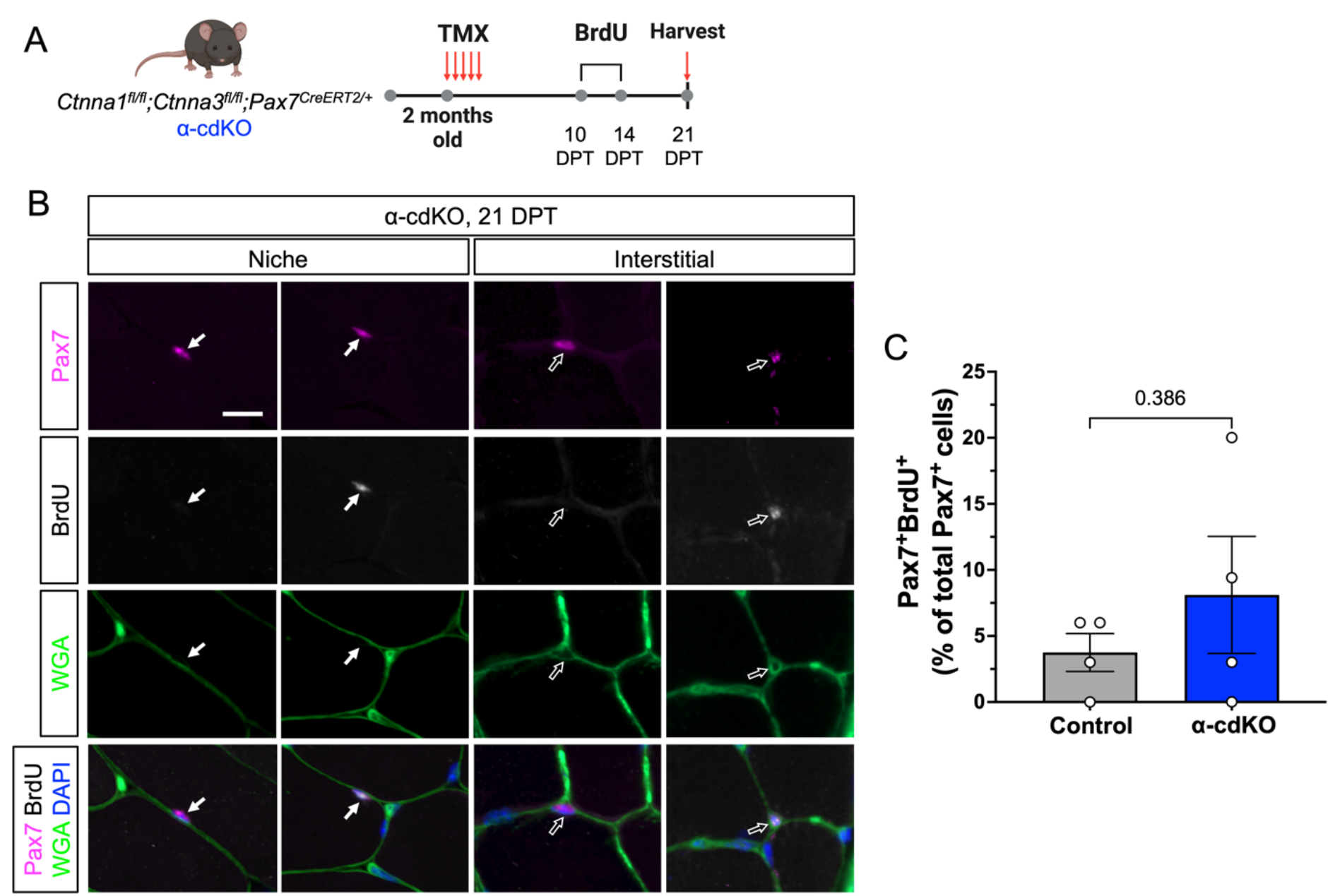
Remaining Pax7^+^ α-cdKO MuSCs are not more likely to incorporate BrdU than controls at 21 days post-tamoxifen. **(A)** Experimental scheme of BrdU pulse-chase experiment to confirm cell cycle progression in MuSCs from control and α-cdKO mice. TA muscles were harvested and snap frozen at 21 DPT for immunofluorescence analysis. Made with BioRender. **(B-C)** Immunostaining of sections through TA muscle from α-cdKO mice 21 DPT (Pax7, magenta; BrdU, grey; WGA, green; and DAPI, blue) (B) showed that there was no significant differences in BrdU incorporation among Pax7^+^ cells (C). Solid arrow denotes cell under the basal lamina; open arrow denotes interstitial cell. Scale bar: 10μm. Each data point represents the average from ten fields (TA muscle sections) from each animal. All data represent n=4 per genotype and represented as mean ± S.E.M with comparisons by two-tailed unpaired Student’s t-test.

**Table S1.**
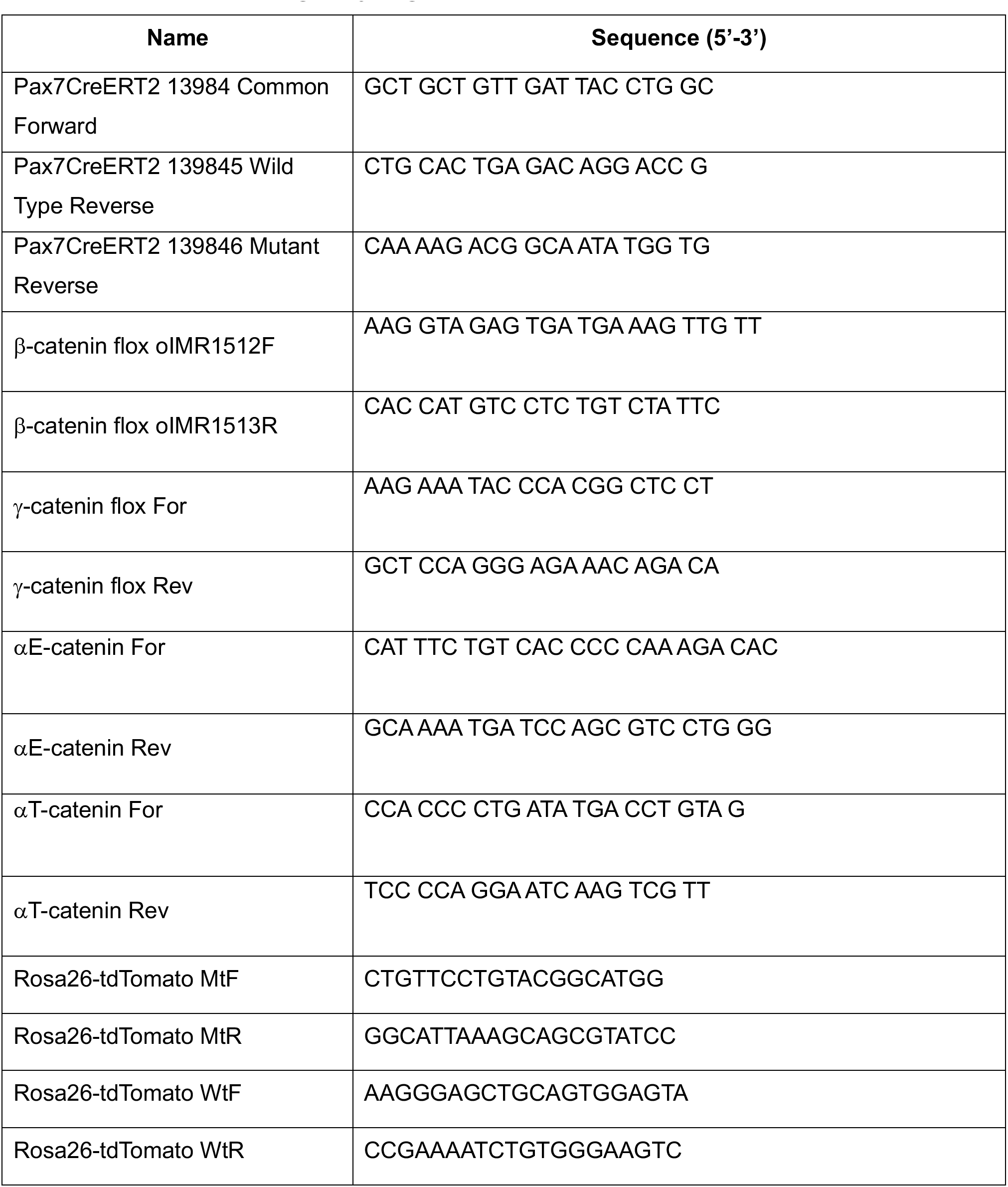
Primers used for genotyping.

**Table S2.** Antibodies and dilutions for immunofluorescence.

| **Antibody** | **Dilution** | **Source** |
| --- | --- | --- |
| Mouse anti-Pax7 | 1:100 | DSHB (Pax7-c) |
| Rabbit anti-Caveolin-1 | 1:200 | Abcam (ab2910) |
| Mouse anti-MyoD | 1:100 | BD Biosciences (554130) |
| Mouse anti-Myogenin | 1:100 | DSHB (F5D) |
| Rabbit anti-Laminin | 1:100 | Sigma (L9393) |
| Mouse anti-β-catenin | 1:200 | BD Biosciences (610154) |
| Rabbit anti-γ-catenin | 1:75 | Cell Signaling Technology (2309S) |
| Rabbit anti-αT-catenin | 1:100 | Proteintech (13974-1-AP) |
| Rabbit anti-αE-catenin | 1:100 | Invitrogen (71-1200) |
| Rabbit anti-M-cadherin | 1:200 | Cell Signaling Technology (40491S) |
| Rat anti-BrdU | 1:300 | Abcam (ab6326) |
| Rabbit anti-Ki67 | 1:500 | Abcam (ab15580) |
| Rabbit anti-cleaved Caspase 3<br>(Asp175) | 1:300 | Cell Signaling Technology (9664) |
| Anti-WGA-488 conjugated | 10 μg/ml | Invitrogen (W11261) |
| Alexa Fluor 647 anti-Mouse IgG1 | 1:500 | ThermoFisher (A-21240) |
| Alexa Fluor 647 anti-Rabbit IgG | 1:500 | ThermoFisher (A-21245) |
| Alexa Fluor 488 anti-Rabbit IgG | 1:500 | ThermoFisher (A-11008) |
| Alexa Fluor 488 anti-Mouse IgG1 | 1:500 | ThermoFisher (A-21121) |
| Alexa Fluor 568 anti-Rat IgG | 1:500 | ThermoFisher (A-11077) |
| Alexa Fluor 568 anti-Rabbit IgG | 1:500 | ThermoFisher (A-11011) |

